# Paracrine role for endothelial IGF-1 receptor in white adipocyte beiging

**DOI:** 10.1101/2021.12.01.470734

**Authors:** Natalie J Haywood, Katherine I Bridge, Cheukyau Luk, Nele Warmke, Katie J Simmons, Michael Drozd, Amy Moran, Sam Straw, Jason L Scragg, Jessica Smith, Sunti Limumpornpetch, Claire H Ozber, Chloe G Wilkinson, Anna Skromna, Natallia Makava, Andrew Walker, Nicole T Watt, Romana Mughal, Kathryn J Griffin, Hema Viswambharan, Nadira Y Yuldasheva, David J Beech, Piruthivi Sukumar, Antonio Vidal-Puig, Klaus K Witte, Stephen B Wheatcroft, Richard M Cubbon, Lee D Roberts, Mark T Kearney

**Affiliations:** Leeds Institute of Cardiovascular and Metabolic Medicine, Faculty of Medicine and Health, University of Leeds, UK.; University of Cambridge Metabolic Research Laboratories, Cambridge, UK; Integrative Vascular Biology Laboratory, Max Delbrück Center for Molecular Medicine in the Helmholtz Association, Berlin, Germany; Department of Optometry and Vision Sciences, University of Huddersfield, UK; Dept of Internal Medicine I, University Clinic, RWTH Aachen University, Aachen, Germany

**Keywords:** Endothelial cell, adipose tissue, IGF-1R, obesity, malonic acid, metabolomics

## Abstract

There are at least two distinct types of thermogenic adipocyte in mammals: a pre-existing form established during development, termed classical brown adipocytes and an inducible form, ‘beige’ adipocytes^1–3^. Various environmental cues can stimulate a process frequently referred to as ‘beiging’ of white adipose tissue (WAT), leading to enhanced thermogenesis and obesity resistance ^4, 5^. Whilst beiging of WAT as a therapeutic goal for obesity and obesity-related complications has attracted much attention^6–9^; therapeutics stimulating beiging without deleterious side-effects remain elusive^10^. The endothelium lines all blood vessels and is therefore in close proximity to all cells. Many studies support the possibility that the endothelium acts as a paracrine organ^11–14^. We explored the potential role of endothelial insulin-like growth factor-1 receptor (IGF-1R) as a paracrine modulator of WAT phenotype. Here we show that a reduction in endothelial IGF-1R expression in the presence of nutrient excess leads to white adipocyte beiging, increases whole-body energy expenditure and enhances insulin sensitivity via a non-cell autonomous paracrine mechanism. We demonstrate that this is mediated by endothelial release of malonic acid, which we show, using prodrug analogues, has potentially therapeutically-relevant properties in the treatment of metabolic disease.

---

Over the past four decades, changes in human lifestyle have contributed to a pandemic of nutritional obesity^15^. In simple terms, obesity occurs due to sustained elevation of calorie intake, most often in the form of lipid and carbohydrate, and/or a decline in energy expenditure^16^. Disruption of this ‘energy balance equation’^17^ can occur at any point in the human life course. In 2015, over 100 million children and 600 million adults were obese worldwide^18^. An unfavourable deviation in the energy balance equation in favour of calorie excess results in ectopic deposition of lipids in tissues such as the liver and skeletal muscle, which are ill-equipped to deal with this challenge. As a result, deleterious perturbations in cellular function lead to type 2 diabetes mellitus, accelerated cardiovascular disease, fatty liver and some cancers (^19^ for review).

Dietary lipids are stored in adipose tissue (AT) of which broadly speaking, there are two types. White AT (WAT) specialised for the storage of energy in the form of triglyceride during nutrient excess undergoes expansive remodelling with adipocytes adopting a hypertrophic/hyperplastic phenotype^20^. The second form of AT is brown AT (BAT)^21^. BAT, unlike WAT, expresses the mitochondrial carrier protein uncoupling protein-1 (UCP-1). UCP-1 uncouples cellular respiration from mitochondrial ATP synthesis, affording BAT the capacity to oxidise lipids and glucose to generate heat^21^.

Recent studies indicate that at least two distinct types of thermogenic adipocyte exist in mammals: a pre-existing form established during development, termed ‘classical brown’, and an inducible form described as ‘beige’^4, 5^. BAT depots, previously thought to be limited to neonates, have also been identified in human adults^1–3^. Beige adipocyte biogenesis can be stimulated by various environmental cues, such as chronic cold exposure, ^4, 5^ in a process frequently referred to as ‘beiging’ of WAT. The potentially favourable metabolic effects of inducing more thermogenic AT in response to nutrient excess, has led investigators to seek new approaches to stimulate WAT beiging.

The insulin/insulin-like growth factor-1 (IGF-1) signalling system evolved millions of years ago to co-ordinate organismal growth and metabolism. During evolution the insulin receptor (IR) and IGF-1 receptor (IGF-1R) diverged from a single receptor in invertebrates, into a more complex system in mammals consisting of the IR, IGF-1R and their respective ligands; insulin, IGF-1 and IGF-II (^22^ for our review). We have previously shown that during calorie excess circulating IGF-1 increases and IGF-1R levels decline, in a range of tissues including the vasculature ^23, 24^.

The endothelium lines all blood vessels, and emerging data support the possibility that the endothelium acts as a paracrine organ^11–14^, including, endothelial to AT signalling^25–27^. Therefore, we explored the role of endothelial IGF-1R as a paracrine modulator in the pathophysiology of obesity, and identified a novel, small molecule mediated beiging mechanism.

## Results

### Murine endothelial IGF-1R knockdown enhances whole-body insulin sensitivity during positive energy balance

To investigate the role of endothelial IGF-1R in the setting of increased energy balance, we generated a tamoxifen-inducible, endothelial cell-specific IGF-1R knockdown mouse (ECIGF-1R^KD^) (Figure 1A-B) with an mTmG reporter to confirm spatially appropriate Cre-recombinase activity (Supplementary figure 1A-C). When unchallenged on a standard laboratory chow diet or challenged with high fat diet (HFD) for two weeks, ECIGF-1R^KD^ mice exhibited no difference in body (Figure 1C) or organ weight (Figure 1D). Glucose tolerance was also unchanged in both chow and HFD fed mice (Figure 1E-F). Insulin sensitivity was similar in chow fed-mice (Figure 1G), but was enhanced in ECIGF-1R^KD^ after HFD feeding for two weeks, compared to wildtype littermates on the same diet (Figure 1H&I). After two weeks HFD, ECIGF-1R^KD^ mice had similar core body temperature (Supplementary Figure 2A) and fasting plasma concentrations of glucose, insulin, IGF-I, free fatty acids, triglycerides, and leptin as wildtype littermates on the same diet (Supplementary Figure 2B-G).

**Figure 1.**
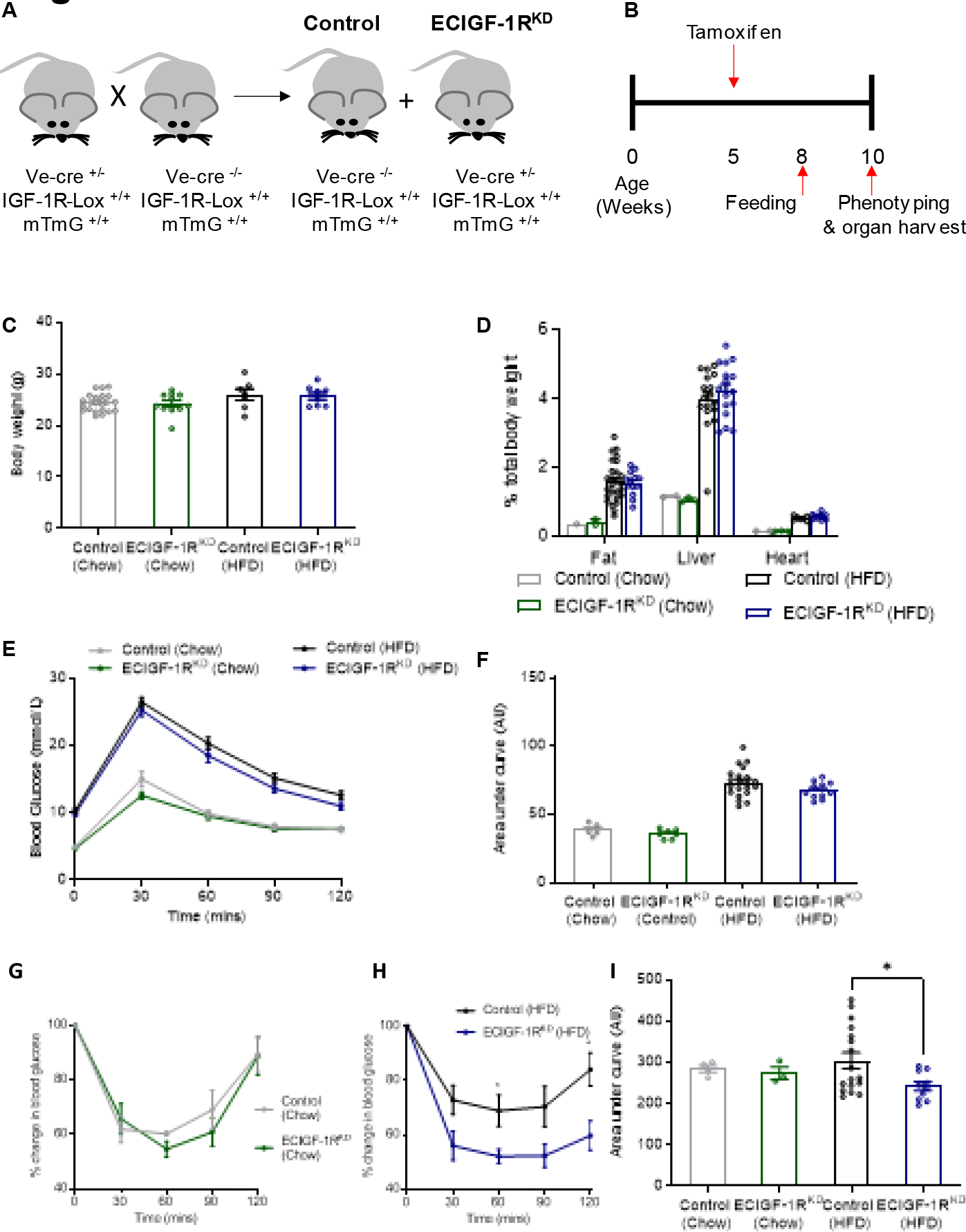
Reduction in murine endothelial IGF-1R expression improves whole body insulin sensitivity in the setting of over nutrition. **A.** Schematic representation of the generation of tamoxifen-inducible endothelial cell specific IGF-1R knock down mice (ECIGF-1R^KD^). **B.** Schematic representation of experimental protocol. **C.** Quantification of body weight from chow-fed control and ECIGF-1R^KD^ mice and from 2-week high fat fed (HFD) control and ECIGF-1R^KD^ mice. (n=Chow 21&11, HFD 7 &8). **D.** Quantification of wet organ weight from chow-fed control and ECIGF-1R^KD^ mice and from 2-week HFD control and ECIGF-1R^KD^ mice (n=Chow 2&3, HFD 26 &16). **E.** Glucose tolerance over time for chow-fed control and ECIGF-1R^KD^ mice and for 2-week HFD control and ECIGF-1R^KD^ mice (n=Chow 5&7, HFD 21&11). **F.** Area under the curve (AUC) analysis for glucose tolerance test for chow fed control and ECIGF-1R^KD^ mice and for 2-week HFD control and ECIGF-1R^KD^ mice (n =Chow 5&7, HFD 21&11). **G.** Insulin tolerance test for chow-fed control and ECIGF-1R^KD^ mice (n =4&3). **H.** Insulin tolerance test for 2-week HFD control and ECIGF-1R^KD^ mice (n =17&10). **I.** The area under the curve analysis for insulin tolerance tests for chow-fed control and ECIGF-1R^KD^ mice and from 2-week HFD control and ECIGF-1R^KD^ mice (n =4,3,17&10). Data shown as mean ± SEM, data points are individual mice. p<0.05 taken as statistically significant using student unpaired two tailed t-test and denoted as *.

**Figure 2.**
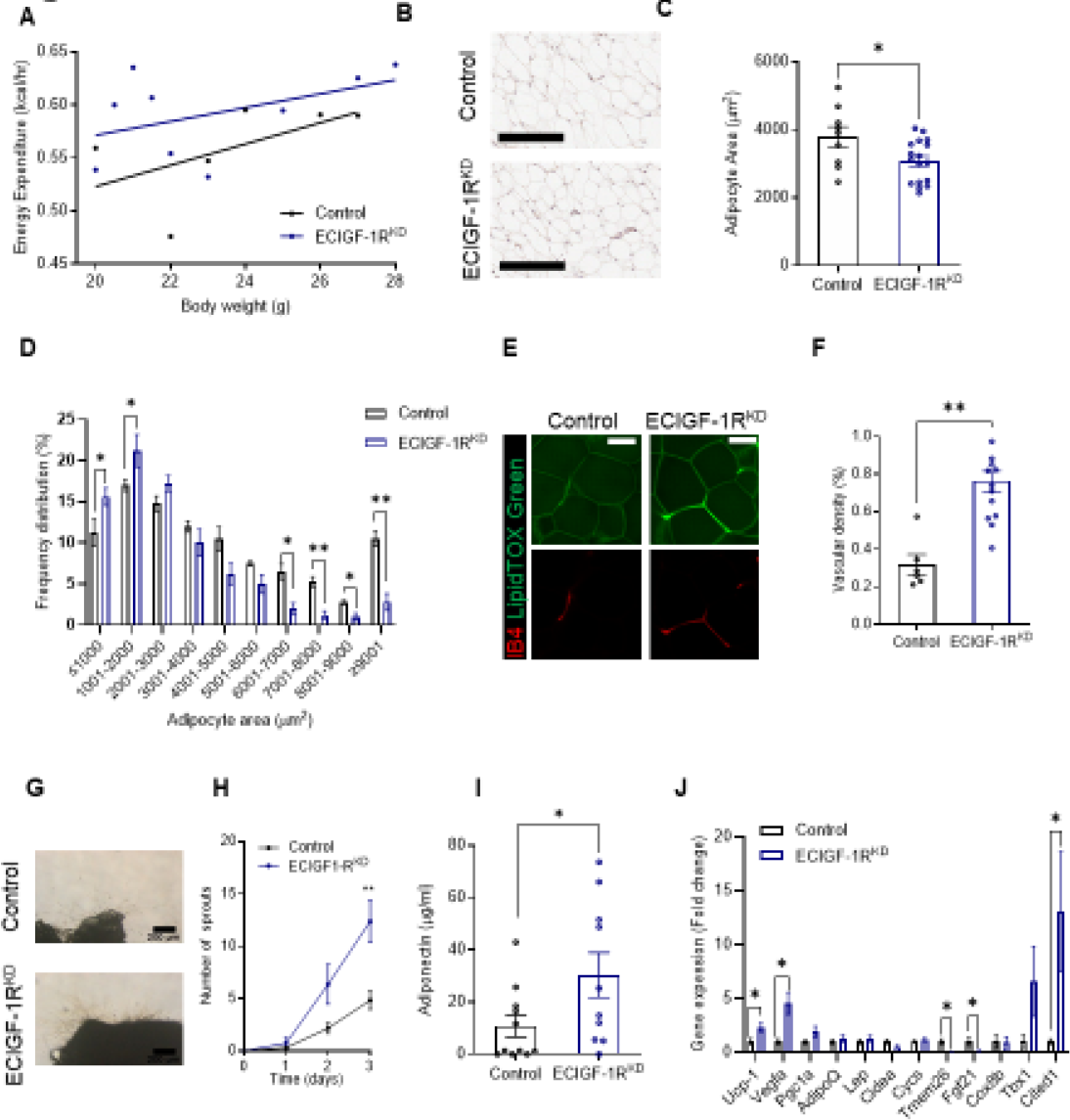
Reduction in murine endothelial IGF-1R expression prevents deleterious remodelling of adipose tissue in the setting of over nutrition. **A.** Energy expenditure in 2-week HFD fed control and ECIGF-1R^KD^ mice (n =6&9). **B.** Representative images of hematoxylin and eosin (H & E) stained white epididymal adipose tissue from 2-week HFD control and ECIGF-1R^KD^ mice (Scale bar = 200µm). **C.** Quantification of adipocyte size from 2-week HFD control and ECIGF-1R^KD^ mice (n =9&17). **D.** Quantification of white epididymal adipocyte size distribution from 2-week HFD control and ECIGF-1R^KD^ mice (n =9&17). **E.** Representative images of isolectin B4 (Red) and LipidTox (Green) stained white epididymal adipose tissue from 2-week HFD control and ECIGF-1R^KD^ mice (Scale bar =100 µm). **F.** Quantification of white epididymal adipose tissue vascularisation from 2-week HFD control and ECIGF-1R^KD^ mice (n =6&14). **G.** Representative images of 2-week HFD control and ECIGF-1R^KD^ white epididymal adipose tissue explants (Scale bar = 200µm). **H.** Quantification of white epididymal adipose tissue neovascularisation from 2-week HFD control and ECIGF-1R^KD^ mice (n =5&5). **I.** Quantification of plasma adiponectin levels from 2-week HFD control and ECIGF-1R^KD^ mice (n =10&11). **J.** Quantitation of white epididymal adipose gene expression from 2-week HFD control and ECIGF-1R^KD^ mice (n =9-17). Data shown as mean ± SEM, data points are individual mice. p<0.05 taken as statistically significant using student unpaired two tailed t-test and denoted as * (**p≤.0.01).

### Endothelial IGF-1R knockdown prevents deleterious remodelling of adipose tissue in the setting of increased energy balance

Historically, WAT was thought to be a simple storage depot for lipid. However, over the past two decades, research has revealed WAT to be a complex and plastic organ (Reviewed ^28^). Targeting AT phenotype to mitigate against the adverse sequelae of obesity has thus received significant attention (Reviewed ^29^). Mechanisms of changing WAT from a storage to thermogenic phenotype has been of particular interest ^6–9^.

After two weeks of HFD, ECIGF-1R^KD^ mice displayed increased energy expenditure in relation to body mass (Figure 2A), with no change in food intake or physical activity (Supplementary Figure 3A-F) compared to wildtype littermates. Epididymal WAT from ECIGF-1R^KD^ mice had smaller adipocytes (Figure 2B-D), increased vascularity (Figure 2E&F) and enhanced *ex vivo* sprouting angiogenesis (Figure 2G&H, Supplementary Figure 4A). This remodelling is contrary to the deleterious phenotype in humans with either a high body mass index or raised HbA1c (Supplementary Figure 5A-I). ECIGF-1R^KD^ mice had similar levels of epididymal WAT fibrosis, lipid accumulation in the liver and interscapular BAT (Supplementary Figure 6A-F). Vascularity in other AT depots (BAT, subcutaneous WAT and perinephric WAT) was unchanged (Supplementary Figure 6G-L), as well as vascularity of other organs including liver and muscle (Supplementary Figure 6M-P), suggesting an epididymal WAT specific effect of reduced endothelial IGF-1R. Chow fed ECIGF-1R^KD^ mice had similar AT vascularity (Supplementary Figure 7A&B), indicative of a specific response to over nutrition. When fed HFD for 2-weeks, ECIGF-1R^KD^ also exhibited higher circulating levels of the beneficial adipokine adiponectin (Figure 2I), an endogenous insulin sensitizer. Epididymal WAT from HFD-fed ECIGF-1R^KD^ also exhibited a change in gene expression including upregulation of *Ucp-1*, *Vegfa* and *Cited1* (Figure 2J), indicative of adipocyte beiging, which was not seen in chow-fed mice (Supplementary Figure 7C).

**Figure 3.**
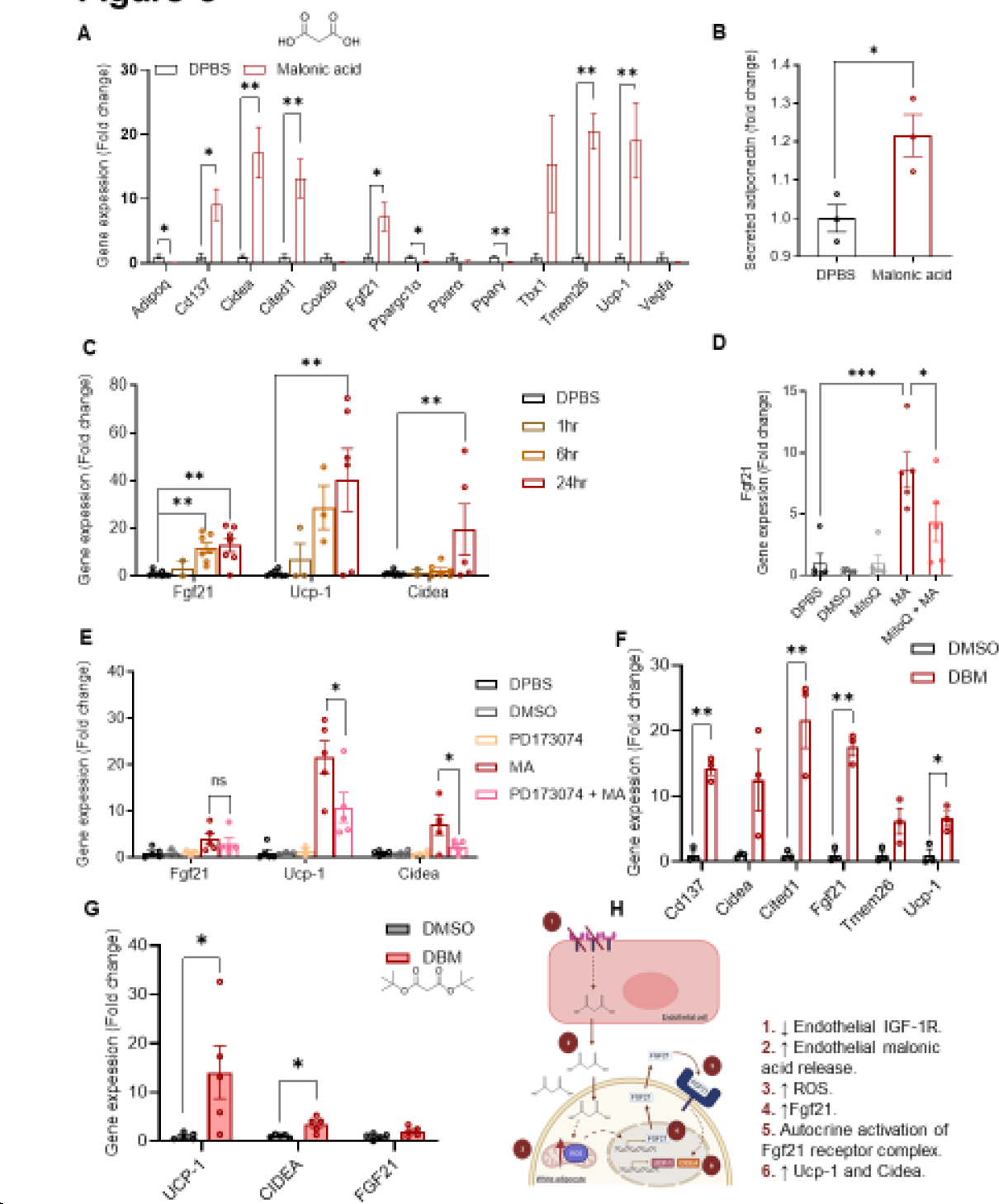
Reduction in murine endothelial IGF-1R expression alters the endothelial secretome and reveals a role for malonic acid in modulating white adipose function. **A.** Quantification of gene expression in 3T3-L1 adipocytes after 24hr 10mM malonic acid stimulation (n =4-7 per treatment). **B.** Quantification of adiponectin secretion in 3T3-L1 adipocytes after 24hr 10mM malonic acid stimulation (n =3 per treatment group). **C.** Quantification of gene expression in 3T3-L1 adipocytes after varying exposure times to 10mM malonic acid stimulation (n =4-7 per treatment group). **D.** Quantification of *Fgf21* gene expression in 3T3 -L1 adipocytes, after treatment with mitoQ and malonic acid 10mM for 24hrs (n =5 per treatment group). **E.** Quantification of *Ffg21, Ucp-1* and *Cidea* gene expression in 3T3-L1 adipocytes after treatment with FGF1R blocker (PD17304) and malonic acid 10mM for 24hrs (n =3-5 per treatment group). **F.** Quantification of gene expression in 3T3-L1 adipocytes after 24hr 10mM di-tert-butyl malonate (DBM) stimulation (n =3 per treatment group). **G.** Quantification of gene expression in human primary adipocytes after 24hr 10mM di-tert-butyl malonate (DBM) stimulation (n =5 per treatment group). **H.** Schematic diagram of the proposed mechanism of EC IGF-1R mediated white adipocyte beiging. Data shown as mean ± SEM, n is an individual experiment. p<0.05 taken statistically significant using student unpaired two tailed t-test or ANOVA and denoted as * (p≤. and is denoted as **).

As discussed above, at least two types of thermogenic adipocyte exist in mammals, BAT and inducible ‘beige’ AT^1–3^. Beige adipocyte biogenesis may be stimulated by various environmental cues ^4, 5^, a process referred to as ‘beiging’. Here we show that a reduction in endothelial IGF-1R expression in the presence of nutrient excess increases beige AT, increases whole-body energy expenditure, and enhances insulin sensitivity.

To explore the chronicity of these findings, a separate cohort of ECIGF-1R^KD^ mice and wildtype littermate controls received HFD for eight weeks. ECIGF-1R^KD^ mice maintained endothelial IGF-1R knockdown (Supplementary Figure 8A&B); body weight, organ weight, core body temperature and glucose tolerance were similar to wildtype littermate controls (Supplementary Figure 8C-H). The enhanced insulin sensitivity of ECIGF-1R^KD^ seen at 2 weeks HFD persisted after 8 weeks of HFD (Supplementary Figure 8I-J). Circulating insulin and IGF-1 concentrations were unchanged (Supplementary Figure 8K-L). Epididymal WAT from ECIGF-1R^KD^ mice after eight weeks HFD still had smaller adipocytes (Supplementary Figure 9A&B). ECIGF-1R^KD^ mice also had reduced lipid accumulation in BAT (Supplementary Figure 9C&D). There was no longer a difference in WAT vascularity (Supplementary Figure 9E&F); however, increased *Vegf* and *Ucp-1* gene expression was retained (Supplementary Figure 9G). There was no difference in WAT collagen deposition (Supplementary Figure 9H-I) or fatty liver in ECIGF-1R^KD^ mice after 8 weeks HFD compared to wildtype littermate controls (Supplementary Figure 9J&K). Taken together, these findings suggest that the advantageous effects of EC IGF-1R knockdown on insulin sensitivity and AT phenotype are retained over longer periods of HFD.

### Endothelial IGF-1R knockdown modifies paracrine modulation of adipocyte function

To probe mechanisms underpinning the favourable changes to WAT in ECIGF-1R^KD^ mice receiving HFD, we investigated the possibility that adipocytes were directly derived from ECIGF-1R^KD^ endothelial cells, as it has been demonstrated that adipocytes of endothelial origin exist in BAT and WAT^30^. However, in our model, Cre activity in ECIGF-1R^KD^ resulted in vascular GFP expression, as intended, but no GFP expressing adipocyte like structures were observed after 2 weeks of feeding, suggesting EC to AT transformation was not occurring (Supplementary Figure 10A).

Since studies continue to emerge supporting the endothelium as a paracrine organ^11–14^, we investigated a potential paracrine mechanism facilitating cross talk between the endothelium and WAT. Treatment of primary human adipocytes with conditioned media from primary EC of ECIGF-1R^KD^ fed HFD for two weeks led to increased *UCP-1*, *CIDEA*, *PGC1a*, *CYCS* and *CD137* gene expression compared to adipocytes cultured in conditioned media from wildtype littermates fed HFD for two weeks (Supplementary Figure 10B). *UCP-1* and *CIDEA* expression induced by ECIGF-1R^KD^ EC conditioned media was preserved following protein denaturation by boiling, suggesting a non-protein signal (Supplementary Figure 10C). We used a metabolomic approach to compare the small molecule secretome of EC from ECIGF-1R^KD^ mice and their littermate controls after two weeks of HFD and found distinct differences in the small molecule endothelial secretome (Supplementary Figure 11A). To our knowledge this is the first study to demonstrate the potential importance of the endothelial small molecule secretome in the pathophysiology of obesity.

### Malonic acid is a novel adipose tissue beiging metabokine

A screen of the upregulated metabolites released from ECIGF-1R^KD^ EC (Supplementary Figure 11B) revealed that malonic acid was sufficient to upregulate *Ucp-1, Cidea, Cd137, Cited1, Fgf21* and *Tmem26* gene expression in 3T3-L1 adipocytes (Figure 3A). A concentration of 10mM malonic (Supplementary Figure 12) increased adiponectin secretion from 3T3-L1 adipocytes (Figure 3B). Time-course experiments demonstrated malonic acid upregulated *Fgf21* gene expression first, followed by *Ucp-1* and *Cidea* gene expression (Figure 3C).

It has previously been shown that malonate, the ionised form of malonic acid, can increase reactive oxygen species (ROS) production^31^. We therefore asked if malonic acid-induced beiging is mediated by ROS. To answer this, we treated adipocytes with MitoQ, a mitochondrial-targeted antioxidant^32^, prior to malonic acid treatment. Removal of mitochondrial-generated ROS by MitoQ attenuated malonic acid-induced *Fgf21* gene upregulation (Figure 3D), demonstrating that malonic acid-induced upregulation of FGF21 signalling is in part dependent on ROS.

FGF21 was previously reported as a paracrine/autocrine beiging mediator in WAT, enriched in murine rosiglitazone-stimulated beige adipocytes and norepinephrine-stimulated brown adipocytes ^33, 34, 35, 36^. An FGF21 receptor blocker (Figure 3E) diminished malonic acid induced upregulation of *Ucp-1* and *Cidea* in 3T3-L1 adipocytes, consistent with previous studies showing FGF21 regulates *Ucp-1* ^37, 38^.

To investigate the therapeutic potential of malonic acid as a beiging agent, we utilized various malonate prodrugs^39^ (Supplementary Figure 13). These malonate prodrugs accelerate malonate delivery *in vivo*^39^. The prodrug, di-tert-butyl malonate (DBM) induced an upregulation of *Cd137*, *Cited1*, *Fgf21* and *Ucp-1* gene expression in 3T3-L1 adipocytes (Figure 3F) and *UCP-1* and *CIDEA* gene expression in human primary adipocytes (Figure 3G), demonstrating a novel therapeutic strategy for inducing beiging in white adipocytes.

Although the action of malonic acid to inhibit succinate dehydrogenase (SDH) was established over 80 years ago^40^, our data demonstrate for the first time that, using an alternative concentration and exposure time, malonic acid leads to beiging of WAT. Mills *et al.,* ^41^ in a shivering thermogenesis model, suggested that elevated succinate led to browning of WAT in a SDH and ROS dependent fashion. In contrast to our findings, Mills *et al*., suggested that malonic acid, by inhibiting SDH, blocked the effect of succinate. However, our data raise the possibility that malonic acid may act via an alternative pathway to induce adipocyte beiging in WAT in a ROS/FGF21 dependent fashion (Figure 3H).

## Summary

In conclusion, our data reveal a hitherto unrecognised non-cell autonomous paracrine mechanism by which a reduction in EC IGF-1R stimulates beiging of WAT in mice challenged by a high-fat diet. Moreover, we present the novel finding that malonic acid, is released by the endothelium when IGF-1R levels are reduced and functions as a ‘metabokine’ leading to beiging of white adipocytes.

## Acknowledgements

We would like to acknowledge the histology service from the Division of Pathology and Data Analytics, Colorectal Pathology Trials, University of Leeds, for sectioning and staining adipose and liver samples. The Faculty of Biological Sciences Bioimaging Facility has received equipment grants from the Wellcome Trust to purchase confocal microscopes used in this project. MTK is the guarantor of this work and, as such, had full access to all the data in the study and takes responsibility for the integrity of the data and the accuracy of the data analysis.

NJH was funded by a British Heart Foundation Project Grant (PG/18/82/34120). CL was funded by a British Heart Foundation PhD studentship (FS/19/59/34896). MD was funded by a British Heart Foundation Clinical Research Training fellowship (FS/18/44/33792). NTW was funded by a British Heart Foundation Project Grant (PG/14/54/30939). LDR was funded by a Diabetes UK RD Lawrence Fellowship (16/0005382) and the Biotechnology and Biological Sciences Research Council (BB/R013500/1). SS is funded by a British Heart Foundation

Clinical Research Training Fellowship (FS/CRTF/20/24071). CHO and RMC were funded by British Heart Foundation Clinical Intermediate Fellowships (FS/12/80/29821). AS was funded by a British Heart foundation Programme grant (RG/15/7/31521). MTK holds a British Heart Foundation Chair in Cardiovascular and Diabetes research, which also funded NM and KJS (CH/13/1/30086).

## Author contribution

NJH, KIB, AS, NM and NYY performed in vivo experiments. NJH, KIB, NW, TS, AV and CHO performed ex vivo experiments. NJH, KIB, CL, KJS, AM, CHW and NTW performed cell culture and in vitro experiments. LDR performed metabolomic analysis. SS, JLS, JS, SL and KW obtained patient samples. NJH, KIB, CL, TS, AV, CHO and KG performed image and data analysis. NJH, MD and MTK wrote the manuscript. DJB, LDR and RMC reviewed the manuscript. DJB, PS, KKW, SBW, RMC and MTK obtained funding.

## Methods

### In vivo animal studies

Mice with tamoxifen-inducible endothelial cell specific knockdown of the IGF-1R receptor (ECIGF-1R^KD^) and their *lox/lox* control littermates, were bred in house from founder animals (VE-Cre #MGI 3848984, Igf1r^(lox)^ #MGI:J:60711^42^, mTmG #MGI:J:124702^43^. Experiments were carried out under the authority of UK Home Office Licence P144DD0D6. Mice were group housed in cages of up to five animals. Only male mice were used for experimental procedures to prevent variability associated with the estrous cycle on adiposity and metabolic readouts ^44, 45^. Cages were maintained in humidity-and temperature-controlled conditions (humidity % at 22°C) with a 2hr light-dark cycle. Genotyping was carried out by Transnetyx commercial genotyping using ear biopsies’. At wee s old, mice were injected with tamo ifen (T5648 Sigma, dissolved in, Corn Oil – also from Sigma, C8267) (1mg/day intra-peritoneal for 5 consecutive days). To induce obesity, 8 week old male mice received high fat diet *ad libitum* for either 2 weeks or 8 weeks (60% of energy from fat) (F1850, Bioserve) with the following composition: protein 20.5%, fat 36% and carbohydrate 36.2% (5.51 kcal/g).

### Insulin and glucose tolerance testing

Mice were fasted overnight prior to glucose tolerance tests or for 2hr prior to insulin tolerance tests. Blood glucose was measured using a handheld Glucose Meter (Accu-Chek Aviva). An intra-peritoneal injection of glucose (1mg/g) or recombinant human insulin (Actrapid; Novo Nordisk) (0.75IU/kg) was given and glucose concentration measured at 30min intervals for 2hrs from the point of glucose/insulin administration. Mice were not restrained between measurements^46^. Data were analysed using GraphPad Prism Area under the curve (AUC) calculations.

### Genotyping of endothelial cell specific knockdown of the IGF-1R receptor (ECIGF-1R^KD^) and their lox/lox control littermates

VE-cre reaction mix;. μl μM orward rimer: ’-GCATTACCGGTCGATGCAACGAGTGATGAG -3’. μl μM e erse rimer: ’-GAGTGAACGAACCTGGTCGAAATCAGTGCG -3’ μl 2 Bio mix red C Master Mix, 3μl water and μl e tracted DNA. C cycle as follows; nitial denaturation 9 °C for min, denaturation 95°C for 15 sec, annealing 51°C for 30 sec, extension 72°C for 1 min and final extension 72°C for 6 min. Denaturation, annealing and extension repeated for 35 cycles. PCR products were then run on a 1.5% agarose gel for 1 hr at 110 V, with a 100 bp ladder. Expected product sizes are Cre Positive – 408 bp.

IGF-1R lo reaction mix; μl μM orward rimer : ’-CTTCCCAGCTTGCTACTCTAG G -3’ μl μM orward rimer2: ’-TGAGACGTAGCGAGATTGCTGTA -3’ μl μM e erse rimer: ’-CAGGCTTGCAATGAGACATGGG -3’ μl 2 Bio mi red C MasterMix, μl water and μl e tracted DNA. PCR cycle as follows; Initial denaturation 94°C for 4 min, denaturation 94°C for 45 sec, annealing 61°C for 45 sec, extension 72°C for 1 min and final extension 72°C for 5 min. Denaturation, annealing and extension repeated for 35 cycles. PCR products were then run on a 1.5% agarose gel for 1 hr at 110 V, with a 100 bp ladder. Expected products sizes are; Wild type – 120 bp, Homozygous – 220 bp and Heterozygous – 120 & 220 bp.

mTm reaction mix;. μl μM Common rimer: ’-CTCTGCTGCCTCCTGGCTTCT-3’ 0.5μl μM Wild type e erse rimer: ’-CGAGGCGGATCACAAGCAATA-3’. μl μM Mutant e erse rimer: ’-TCAATGGGCGGGGGTCGTT-3’ μl 2 Bio mi red C Master Mix, 2. μl water and μl e tracted DNA. C cycle as follows; nitial denaturation 94°C for 2 min, denaturation 94°C for 30 sec, annealing 62°C for 30 sec, extension 72°C for 30 sec and final extension 72°C for 10 min. Denaturation, annealing and extension repeated for 35 cycles. PCR products were then run on a 1.5% agarose gel for 1 hr at 110 V, with a 100 bp ladder. Expected product sizes are; Wild type -330 bp, Homozygous -250 bp, and Heterozygous -250 & 350 bp.

### Confirmation of tamoxifen induction of mT to mG

Founder mTmG mice were obtained from the Jackson Laboratory (Bar Harbor, ME, USA). In the absence of Cre recombinase, mTmG mice constitutively express mTdTomato, a non-oligomerizing DsRed variant. After tamoxifen induction and therefore following exposure to Cre recombinase and excision of the mTdTomato expression cassette, the rearranged mTmG transgene converts to the expression of mGFP (green fluorescent protein). Both mTdTomato and mGFP are membrane-targeted, allowing for delineation of single cells using fluorescence microscopy. Mice were perfuse-fixed with 4% paraformaldehyde (PFA). Femoral arteries were excised, permeabilised (0.1% TritonX-100 in PBS) and blocked (Serum free protein block, DAKO), before overnight incubation with a rabbit polyclonal antibody to mouse CD31 (ab28364, Abcam) followed by overnight incubation with a goat polyclonal anti-rabbit conjugated to Chromeo642 (ab60319, Abcam, UK). Arteries were then mounted *en face* on slides using DAPI (DAPI-Fluoromount-G, Southern Biotech) to define nuclei. Confocal microscopy (LSM 700, Zeiss, UK) was used to define CD31, mGFP and mTdTomato fluorescence.

### Primary endothelial cell isolation and culture

Primary endothelial cells (PECs) were isolated from lungs, as previously reported ^47, 48^. Briefly, lungs were har ested, washed, finely minced, and digested in Han s’ balanced salt solution containing 0.18 units/mL collagenase (10 mg/mL; Roche) for 45min at 37°C. The digested tissue was filtered through a 70-μm cell strainer and centrifuged at, 1000 RpM for min. The cell pellet was washed with PBS/0.5% BSA, centrifuged, re-suspended in 1mL PBS/0.5% BSA, and incubated with 1×10^6^ CD146 antibody–coated beads (Miltenyi Biotech, 130-092-007) at 4°C for 30min. Bead-bound PEC were separated from non–bead-bound cells using a magnet. Cells were re-suspended in 2ml supplemented endothelial growth medium–MV2 (PromoCell) and seeded on a 6 well fibronectin coated plates. Cells were cultured at 37°C in 5% CO2 with twice-weekly media changes until confluent.

### Quantification of protein expression

Cells were lysed or tissue mechanically homogenised in lysis buffer (Extraction buffer, FNN0011) and protein content was quantified by a BCA assay (Sigma-Aldrich, St. Louis, MO). Twenty micrograms of protein were resolved on a 4-12% Bis-Tris gel (Bio-Rad, Hertfordshire, UK) and transferred to nitrocellulose membranes. Membranes were probed with antibodies diluted in 5% BSA (IGF-1R and IR, Cell signalling #9750 and #3025 respectively), before incubation with appropriate secondary horseradish peroxidase-conjugated antibody. Blots were visualised with Immobilon Western Chemiluminescence HRP Substrate (Merck Millipore, Hertfordshire, UK) and imaged with Syngene chemiluminescence imaging system (SynGene, Cambridge, UK). Densitometry was performed in ImageJ.

### Plasma samples

Fasting plasma samples were collected from the lateral saphenous vein (EDTA collection tubes Sarstedt 16.444) and spun at 10,000 RPM for 10min in a bench top centrifuge. Fasting plasma insulin (90080, CrystalChem), IGF-I (MG100, R and D systems), leptin (EZML-82K,Merk-Millipore), adiponectin (EZMADP-60K, Merk-Millipore) triglycerides (Abcam Ab65336) and free fatty acids (Abcam, ab65341) were measured as per manufactures instructions.

### Metabolic phenotyping

Metabolic parameters were measured by indirect calorimetry using Comprehensive Lab Animal Monitoring Systems (CLAMS)(Columbus Instruments). In brief, mice were individually housed for 5 days and measurement of their oxygen consumption, carbon dioxide production, food intake, and locomotor activity were continuously recorded^49^. For each mouse, a full 24 hour period, taking into account sleep and wake cycles, was analysed after an acclimatisation period ^50^. Core body temperature was measured using a rectal temperature probe (Vevo2100 (Visualsonics, FujuFilm) with an Indus rectal temperature probe).

### Murine tissue samples

After either 2 or 8 weeks of high fat feeding, all mice were sacrificed using terminal anaesthesia and organ weights measured using a standard laboratory balance.

### Quantification of gene expression

RNA was isolated from tissue and cells samples (NEB, T2010S). The concentration of RNA in each sample (ng/ul) was measured using a Nanodrop. cDNA was reverse transcribed from the RNA samples (NEB, E3010L). Quantitative PCR (qPCR) was performed using a Roche LightCycler 480 Instrument II, using SYBR Green PCR Master Mix (Bio-Rad, 1725120) and relevant primers (Table 1). The ‘cycles to threshold’ (cT) was measured for each well, the average of triplicate readings for each sample taken, normalised to GAPDH, and finally the differential expression of each gene was calculated for each sample.

**Table 1:**
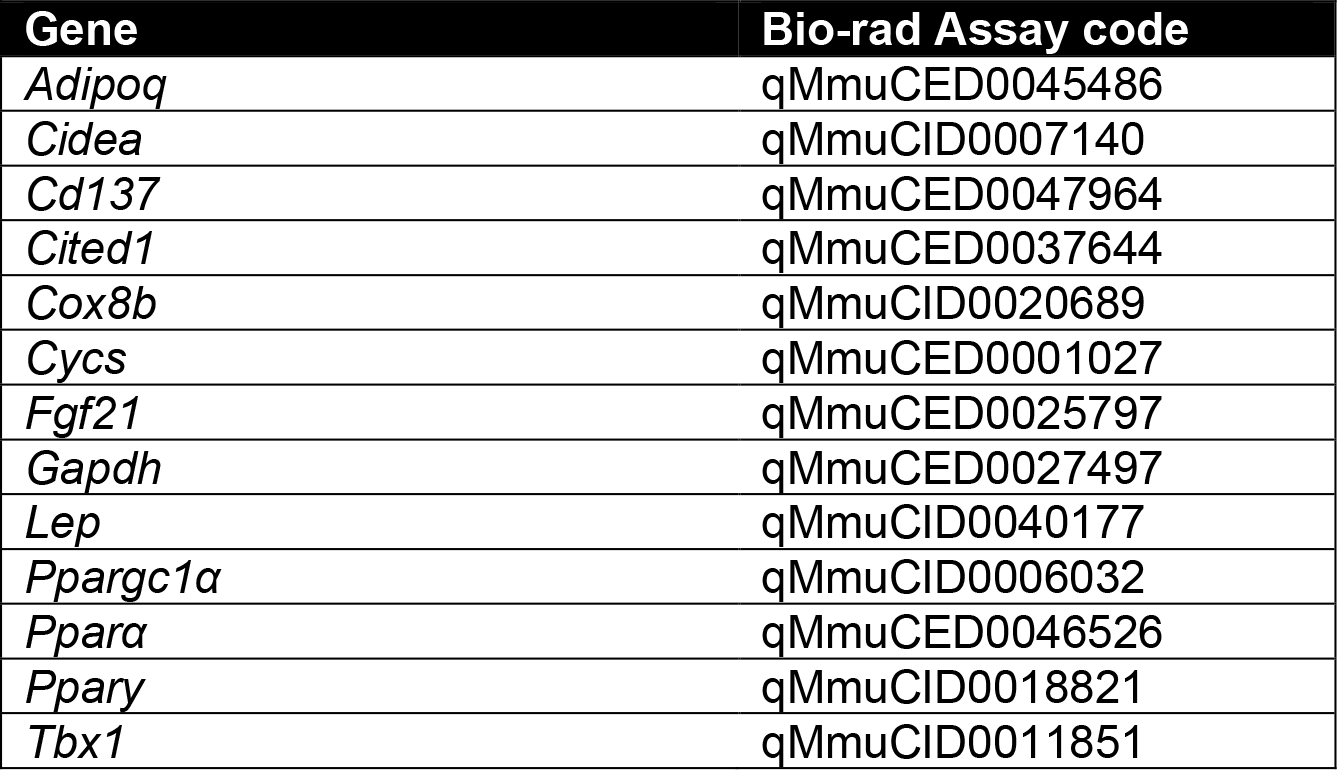
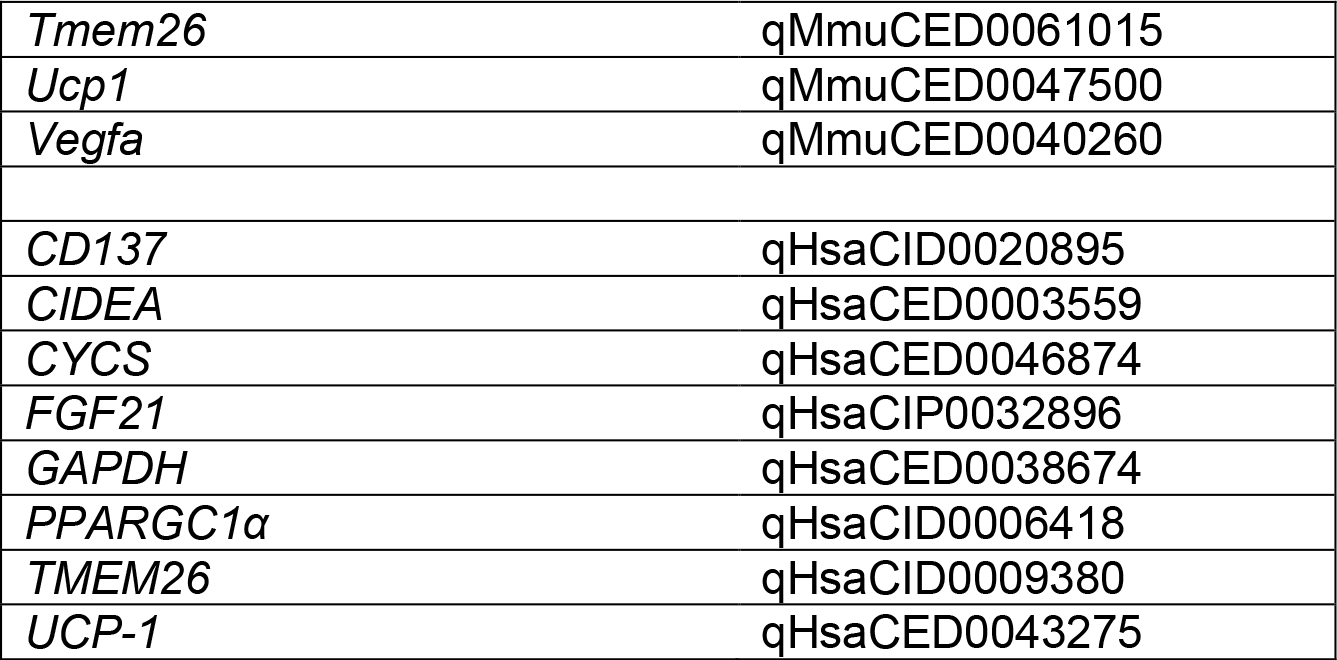
Primers for qPCR.

### Histological assessment of adipocyte size, fibrous tissue and non-alcoholic fatty liver disease

Samples for histology were fixed in 4% PFA for at least 24hrs and then processed into paraffin blocks. 5µm sections were taken, slides were stained with haematoxylin and eosin to assess gross morphology or Picro-sirius red for collagen deposition. Slides were imaged using an Olympus BX41 microscope at 10x and 20x magnification. For assessment of adipocyte size, three separate fields of view for each sample were assessed. For each one, the average was taken of 20 randomly selected independent cells measured using ImageJ. For collagen deposition, the percentage of the sample staining positive for collagen was measured using thresholding in ImageJ, and again was taken as the average in at least three independent areas of the sample. For assessment of non-alcoholic fatty liver disease (NAFLD) in sections of murine liver, a validated rodent NAFLD scoring system was used^51^, which takes into account micro and macro-steatosis, inflammation and hypertrophy. Each sample was assessed by at least two blinded independent verifiers (NH, KB or NW), and the average score per sample taken.

### Quantification of tissue vascularity

Adipose tissue was fixed in 1% paraformaldehyde (PFA), and allowed to fix for 2hrs at room temperature; samples were transferred into phosphate buffered saline (PBS) for longer storage. Samples were incubated overnight with lectin from *Ulex europaeus* Alexa Fluor 594 (73873, Sigma) (For human samples) or Isolectin B4 Alexa Fluor 647 (I32450, Thermo Fisher Scientific) (for murine samples), diluted 1:100 in 5% BSA in PBS at 4°C. After washing with PBS, they were incubated with HCS LipidTOX (H34475, Thermo Fisher Scientific) diluted 1:200 in PBS for 20mins at room temperature. Whole tissue sections were then mounted onto slides beneath cover slips using a silicone spacer (Grace bio-labs, 664113), with Prolong Gold (P36930, Thermo Fisher Scientific). Vascular density (the proportion of each image stained with lectin) was measured using thresholding in ImageJ. Green staining is a composite of GFP and Lipidtox in mice samples, green is not quantified.

Organs (Muscle and liver) were harvested under terminal anaesthesia and fixed in 4% PFA for 1hr at room temperature. Organs were then embedded in Optimal Cutting Temperature compound (OCT) (Cellpath, KMA-0100-00A) and stored at -80 until sectioned. 10µM sections were taken using a Leica CM3050 S Research Cryostat. Slides were blocked and permeabilised in PBS + 0.25% Triton-X100 + 1% BSA + for one hour, then stained with Isolectin B4-Alexa Fluor-488 (Invitrogen I21411) at 1/100 in PBS + 0.25% Triton + 1% BSA for 1hr. Slides were washed three times in PBS and mounted with a coverslip using Prolong Gold with DAPI (P36931, ThermoFisher). Slides were then imaged using laser scanning confocal microscopy (LSM880, Zeiss), with 8 areas of each sample imaged. Vascular density (the proportion of each image stained with IB4) was measured using thresholding in ImageJ.

### Assessment of neovascularisation in white adipose tissue

Angiogenesis assays from adipose tissue were performed using a modified technique based on previously published methods ^52^. In sterile conditions, any surface blood vessels were dissected from the adipose tissue sample, before it was cut into pieces no bigger than 1mm^3^. For each sample, at least 20 sections were embedded into a fibrin matrix. The fibrin matrix was achieved by combining 12.5ul of 50 U/ml thrombin (Sigma-Aldrich T-3399) with 500ul of a mix containing 4 U/ml aprotinin (Sigma-Aldrich A-1153) and 2 mg/ml fibrinogen type 1 (Sigma-Aldrich F-8630), and adding a piece of adipose tissue into the well before the matrix had set. The plates were then incubated at room temperature for 20 minutes, and then at 37°C for a further 20 minutes to ensure that the matrix had fully formed around the piece of adipose tissue. One millilitre of media was then carefully pipetted onto the top of each well, and plates were cultured at 37°C, 5% CO2 for up to 7 days. The media was discarded and replaced every other day throughout the culture period. Each day, the samples were imaged at 4x magnification on Olympus florescent microscope CKX41 and number of endothelial sprouts coming from each piece of fat was counted. For each sample, the average number of sprouts per section was calculated, as well as the number of sections which had sprouted.

### Human adipose tissue explants

Human subcutaneous white adipose tissue was obtained after informed consent from patients undergoing pacemaker implantation at Leeds Teaching Hospitals NHS Trust, Leeds, United Kingdom, after ethical approval (REC: 11/YH/0291). Adipose tissue was removed from the area between the skin and pectoralis major, under local anaesthetic (1% lidocaine). Patient demographics are provided in table 2.

**Table 2.** Patient characteristics.

|  | Mean (±SEM) |
| --- | --- |
| Male | 28 |
| Female | 15 |
| Diabetes | 12 |
| Age (Years) | 68 (1.98) |
| Weight (Kg) | 86.23 (3.12) |
| Height (m) | 1.69 (0.013) |
| BMI (Kg/m<sup>2</sup>) | 29.65 (0.79) |
| HbA1c (mmol/mol) | 46.75(2.469) |

### Conditioning media

When PECS reached confluency, supplemented growth media was removed and replaced with basal endothelial growth medium–MV2 (Promocell) for 24hrs. Conditioned media was then removed and used in further experiments as described.

### Quantification of browning in human adipocytes

Human primary adipocytes (PromoCell, C-12730) were seeded (10,000 cells/cm^2^) in 24 well plates (Costar, Corning, NY, USA) and grown until confluence (37°C, 5% CO2) in PromoCell Preadipocyte Growth Medium (C-27410, 0.05 mL/mL fetal calf serum, 0.004 mL/mL endothelial cell growth supplement, 10 ng/mL epidermal growth factor, 1 µg/mL hydrocortisone, 90 µg/mL heparin) as previously described^49^. To differentiate confluent pre-adipocytes, growth medium was replaced by PromoCell Adipocyte Differentiation Medium (C-27436, 8 µg/mL d-Biotin, 0.5 µg/mL insulin, 400 ng/mL dexamethasone, 44 µg/mL IBMX, 9 ng/mL L-thyroxine, 3 µg/ml ciglitazone) for 48 hours (day 0). Differentiation medium was subsequently replaced (day 2) with PromoCell Adipocyte Nutrition Medium (C-27438, 0.03 mL/mL fetal calf serum, 8 µg/mL d-Biotin, 0.5 µg/mL insulin, 400 ng/mL dexamethasone) for the remainder of the differentiation period (up to day 14). All Cell medium was supplemented with 1% penicillin-streptomycin (10,000 units/mL penicillin, 10 mg/mL streptomycin). Conditioned media was then added to the differentiated human adipocytes for 24hrs before the cells were lysed and RNA extracted for gene expression analysis. To determine if browning was caused by a protein or small molecule, parallel experiments were conducted whereby the conditioned media was boiled for 5mins at 95°C, to denature any proteins, before being added to the human adipocytes.

### Metabolite & pro-drug screening

Mouse 3T3-L1 preadipocytes were cultured in 10% (v/v) CS/DMEM containing 4.5 g/l glucose and 1mM Sodium Pyruvate and supplemented with 1XAntibiotic Antimycotic Solution and incubated at 37°C in 5% CO2 for two days upon splitting. Two days after splitting, the media was replaced by 10% (v/v) FBS/DMEM to grow the cells to confluency. After two days of post-confluency (equivalence of day 0), adipocyte differentiation was initiated with MDI induction media (10% FBS/DMEM, 0.5mM IBMX, 1µM dexamethasone and 1µg/mL insulin). On day 2, the MDI induction media was replaced by insulin media (10% FBS/DMEM supplemented with 1µg/mL insulin). From day 4 onwards, the media was replaced by 10% FBS/DMEM every two days. Full differentiation was achieved between day 7 and day 10. Mature adipocytes were subjected to metabolite stimulation. Metabolites, Malonic acid^31^ or pro-drugs (Table 3) and their solvents, DPBS or DMSO, were applied for 24 hours at 37°C in 5% CO2.

**Table 3:**
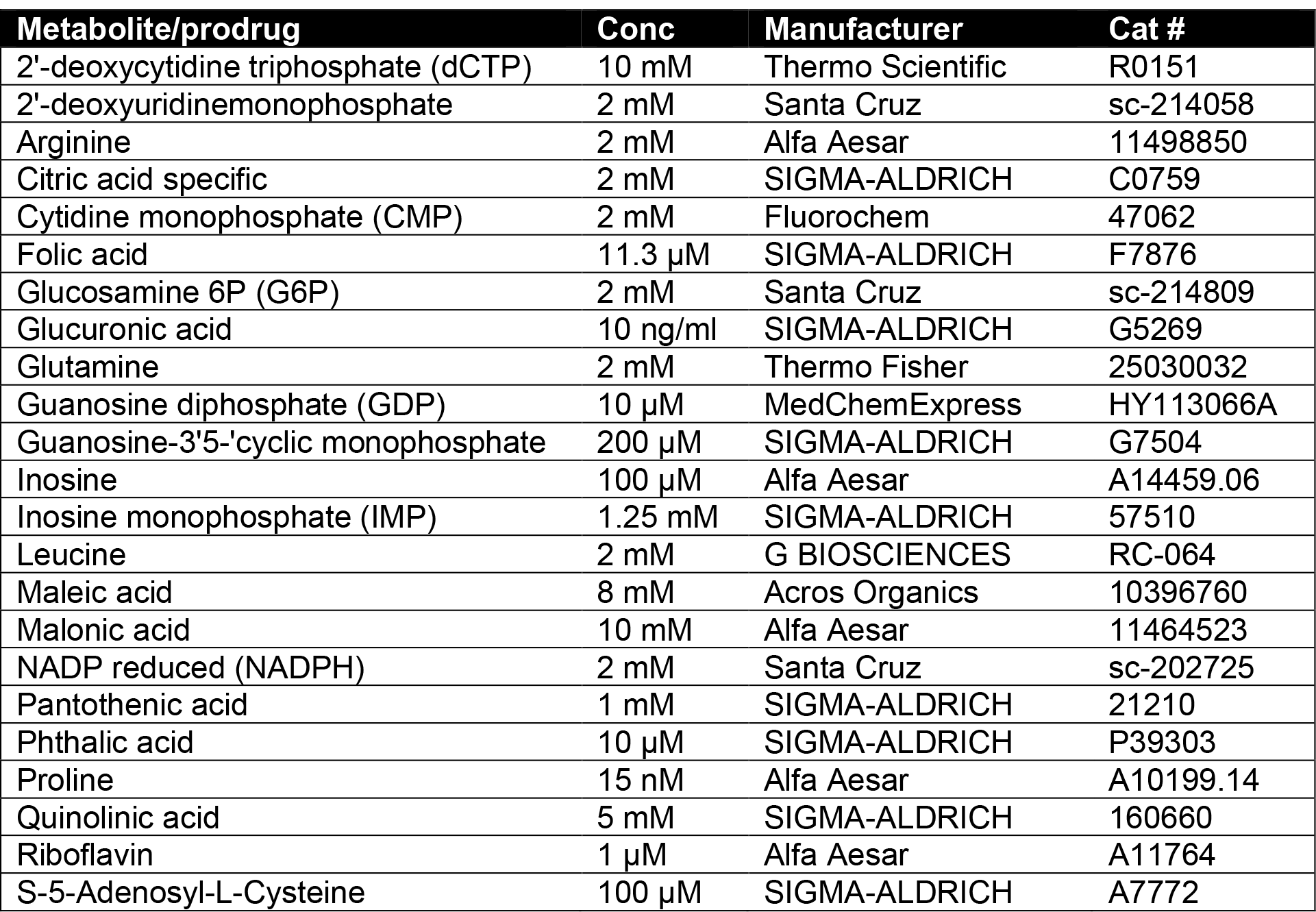
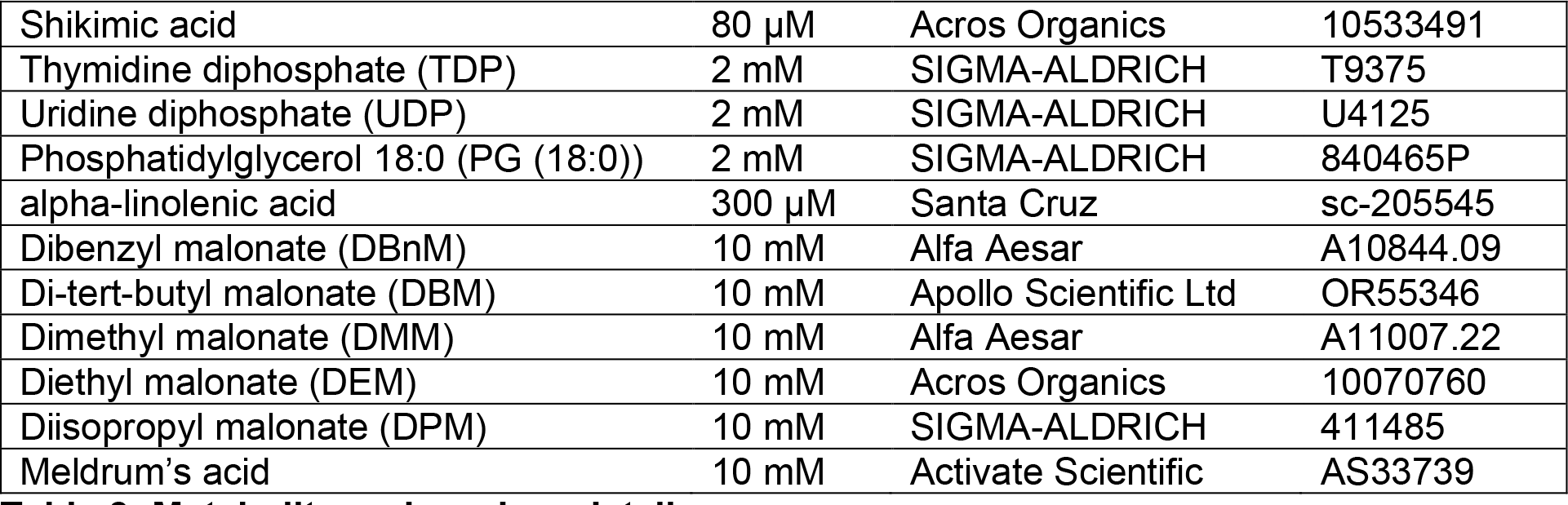
Metabolite and prodrug details.

### Adiponectin secretion

After stimulation of the mature 3T3-L1 adipocytes, the conditioned media was collected and centrifuged at 13,400 rpm for 10 min at 4°C to pellet cell debris. The supernatants were then used to quantify the level of adiponectin using Mouse Adiponectin ELISA kit (Merck Millipore #EZMADP-6 K) according to the manufacturer’s instructions.

### FGFR1 blocker

20nM PD173074 (Apexbio #A8253) was applied an hour prior to 24-hour malonic acid treatment ^53^.

### Mitochondria-targeted antioxidant

100nM Mitoquinone (MitoQ; MedChemExpress LLC #HY-100116A) was applied 30min prior to 24-hour malonic acid treatment ^41^.

### Separation of conditioned media into aqueous and lipid fractions

600µl of 2:1 methanol:chloroform was added to 1 ml of conditioned media, followed by 200 µl of water and an additional 200µl of chloroform, vortexed and then centrifuged at 13.1g for 20mins. The top layer (aqueous layer containing the aqueous metabolites) was pipetted off and placed into the evacuation centrifuge at 40°C for 6 hours. The protein disc (middle layer) was discarded and the final bottom layer (containing lipid metabolites) was transferred into a clean Eppendorf and left overnight in a fume hood at room temperature until all chloroform had evaporated. Both the lipid and aqueous metabolites were stored at -80°C.

### Aqueous Sample Preparation

Samples were reconstituted in 1 ml sample resuspension buffer (95% acetonitrile and 5 % mobile phase A). Mobile phase A = 95% water, 5% acetonitrile, 20mM ammonium acetate and 20mM ammonium hydroxide, pH = 9. Samples were vortex mixed and the extracted metabolites were transferred to a 2 mL glass vial.

### Liquid Chromatography

A SC EX E ion C™ AD H C system with a una 3 µm NH2 Å, 4.6 mm column (Phenomenex) was used. Mobile phase A = 95% water, 5% acetonitrile, 20mM ammonium acetate and 20mM ammonium hydroxide, pH = 9; Mobile phase B = 95% acetonitrile and 5% mobile phase A and 20 mM ammonium hydroxide. The flow rate was set at 350 µL/min. The wash solvent for the autosampler was 20/20/60 methanol/acetonitrile/isopropanol. The injection volume was 2 µL, and the column was kept at 40°C. The gradient method was 100% B for 2 minutes, then to 85% B for 3 minutes, then to 30% for 10min, then to 2% B for 5min, then 100% for 10min.

### Mass Spectrometry

A SCIEX QTRAP® 6500+ with IonDrive Turbo V source was used. MS source parameters are Curtain Gas was 30 for both (+) and (-). Collision Gas was high for both (+) and (-), Ionspray voltage was 5500 for (+) and -4500 for (-). Temperature was 500 for both (+) and (-), Ion source gas 1 was 35 for both (+) and (-), ion source gas was 45 for both (+) and (-), delustering potential was 93 for (+) and -93 for (-), entrance potential was 10 for (+) and -10 for (-) and collision cell exit potential was 10 for (+) and -10 for (-).

### Quantification and statistical analysis

Priori sample size calculations for animal experiments were performed using our published pilot data using the online software package from Vanderbilt University for multiple types of power analysis (https://biostat.app.vumc.org/wiki/Main/PowerSampleSize). All data are shown as mean ± SEM. Individual mice or replicates are shown as individual data points. All image analysis was performed in ImageJ. earsons’ correlation coefficients (*r*) were calculated to assess the link and the degree of relation between BMI and HbA1C and various fat markers. Student 2-tailed unpaired t-test or one-way ANOVA (where appropriate) were used for statistical analyses and were performed with GraphPad Prism software version 7. * denotes P ≤0.05 and ** P ≤0.01. Exact details can be found in figure legends.

### Data availability

The data that support the findings of this study are available from the corresponding author upon reasonable request.

---

**Supplementary figure 1.**
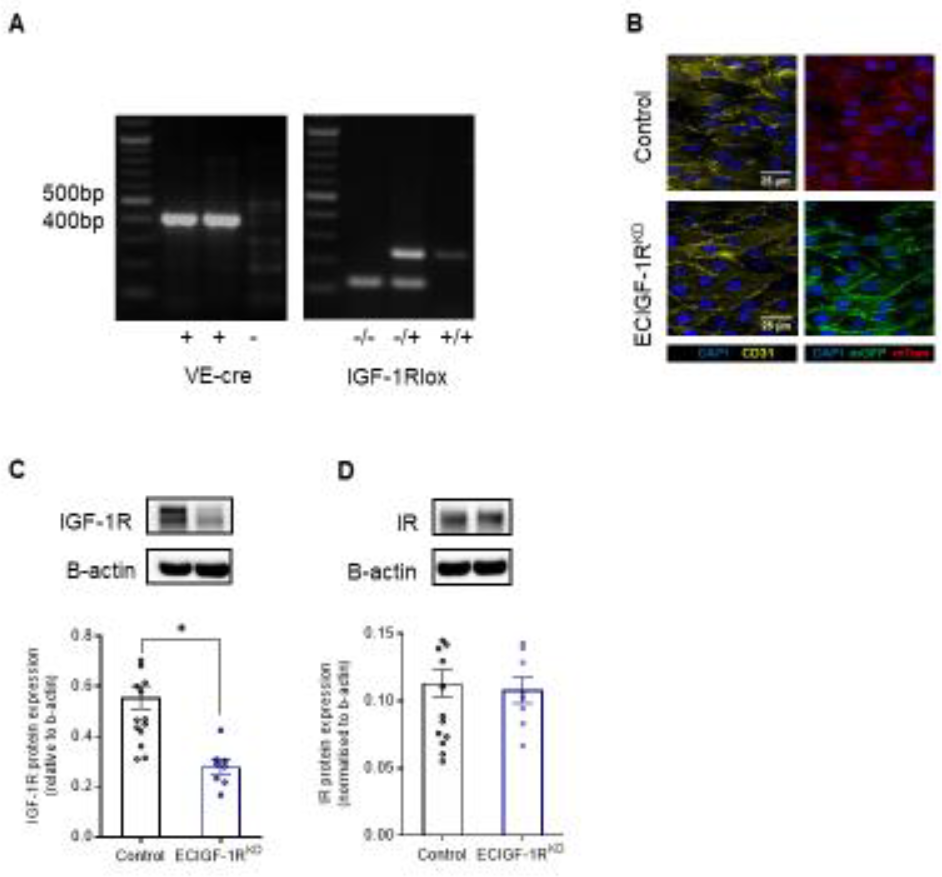
Confirmation of IGF-1R reduction in murine model of endothelial specific IGF-1R knockdown. **A.** Representative images of genotyping during the generation of ECIGF-1R^KD^. **B.** Representative images of *en face* stained femoral arteries from tamoxifen-induced ECIGF-1R^KD^ mice and control littermates confirming a switch from mT to mG with tamoxifen (Scale bar = 25µm). **C.** Quantitation of primary murine endothelial cell expression of IGF-1R from 2-week HFD control and ECIGF-1R^KD^ mice (n =16&8). **D.** Quantitation of primary murine endothelial cell expression of insulin receptor (IR) from 2-week HFD control and ECIGF-1R^KD^ mice (n =16&8). Data shown as mean ± SEM, data points are individual mice. p<0.05 taken as statistically significant using student unpaired two tailed t-test and denoted as *.

**Supplementary figure 2.**
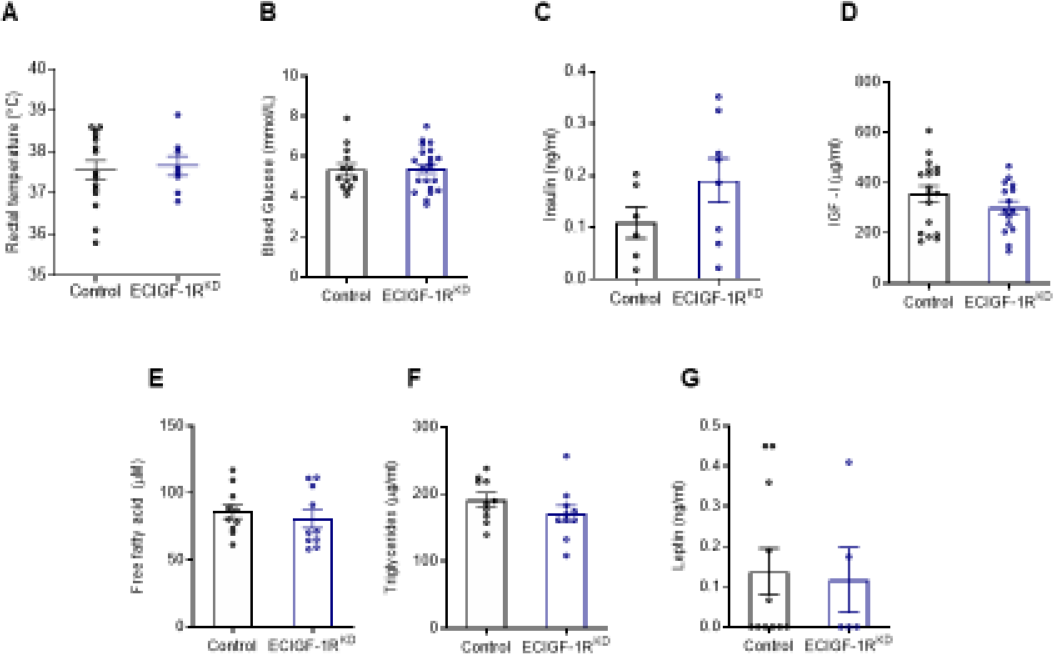
No difference in metabolic plasma markers from mice with endothelial specific IGF-1R reduction in the setting of over nutrition. **A.** Core body temperature of 2-week HFD control and ECIGF-1R^KD^ mice (n =15&8). **B.** Fasting blood glucose levels of 2-week HFD control and ECIGF-1R^KD^ mice (n =13&21). **C.** Fasting plasma insulin levels from 2-week HFD control and ECIGF-1R^KD^ mice (n =6&8). **D.** Fasting circulating plasma IGF-1 levels from 2-week HFD control and ECIGF-1R^KD^ mice (n =18&15). **E.** Fasting plasma free fatty acids levels from 2-week HFD control and ECIGF-1R^KD^ mice (n =10&10). **F.** Fasting plasma triglyceride levels from 2-week HFD control and ECIGF-1R^KD^ mice (n =10&10). **G.** Fasting plasma leptin levels of 2-week HFD control and ECIGF-1R^KD^ mice (n =11&5). Data shown as mean ± SEM, data points are individual mice. p<0.05 taken as being statistically significant using student unpaired two tailed t-test and denoted as *.

**Supplementary figure 3.**
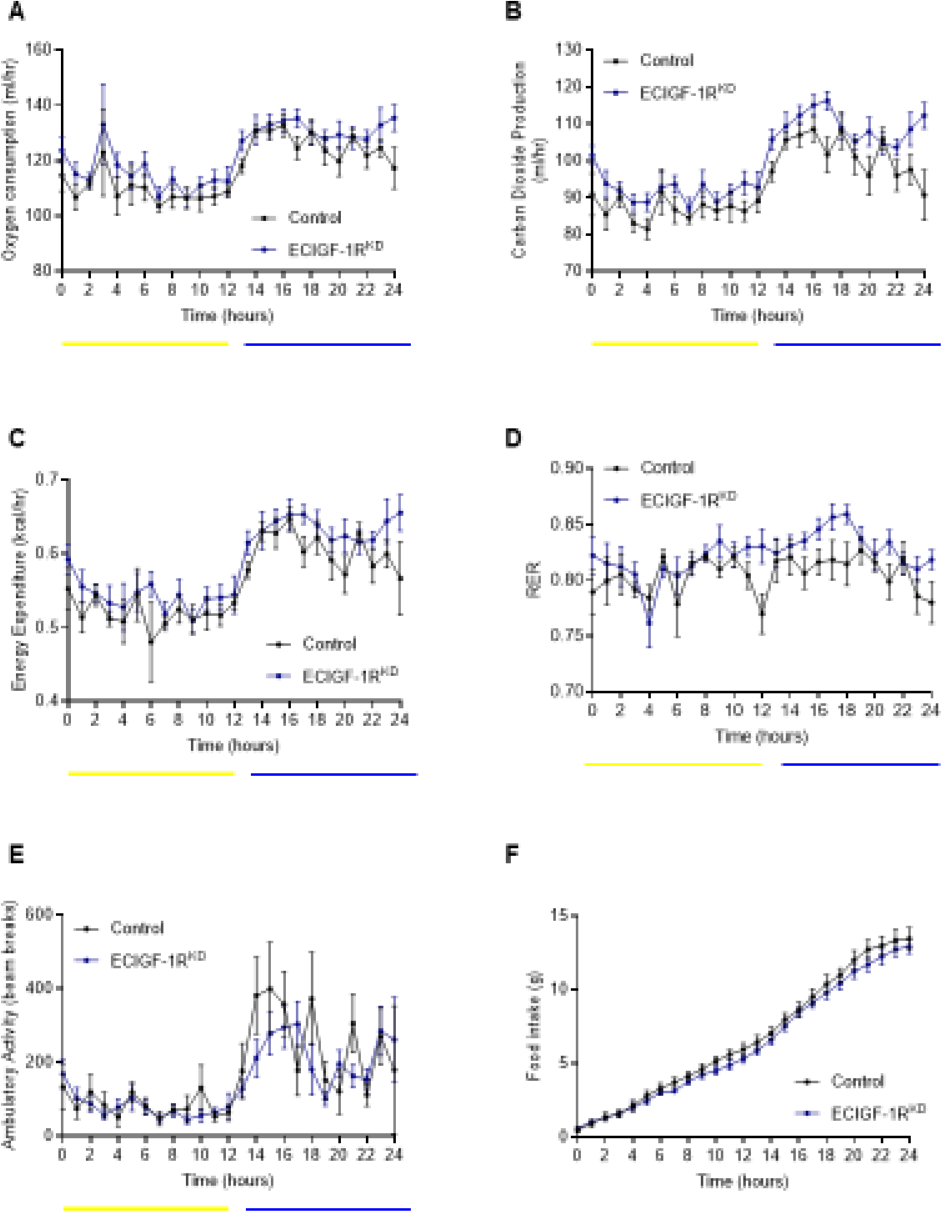
Characterisation of energy expenditure in mice with reduced endothelial IGF-1R expression after 2 weeks of high fat diet. **A.** Oxygen consumption for 2-week HFD control and ECIGF-1R^KD^ mice (n= 6&9). **B.** Carbon dioxide production for 2-week HFD control and ECIGF-1R^KD^ mice (n= 6&9). **C.** Energy expenditure for 2-week HFD control and ECIGF-1R^KD^ mice (n= 6&9). **D.** Respiratory exchange ratio for 2-week HFD control and ECIGF-1R^KD^ mice (n= 6&9). **E.** Activity levels for 2-week HFD control and ECIGF-1R^KD^ mice (n= 6&9). **F.** Cumulative food consumption for 2-week HFD control and ECIGF-1R^KD^ mice (n= 6&9). The light/dark cycle for graphs A-E are shown as follows; Light in yellow and dark in blue. Data shown as mean ± SEM, p<0.05 taken as being statistically significant using student t-test and denoted as *. Metabolic parameters were measured by indirect calorimetry, ANOVA testing was performed using mass as a co-variant (ANCOVA testing) using calrapp.org.

**Supplementary figure 4.**
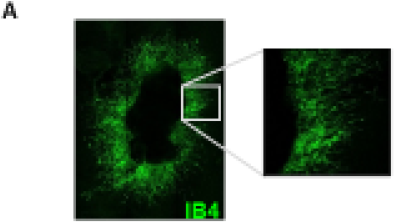
Confirming endothelial cells in neovascularisation. **A.** Staining of adipose tissue explants confirms sprouts are endothelial with positive isolectin B4 staining (green).

**Supplementary figure 5.**
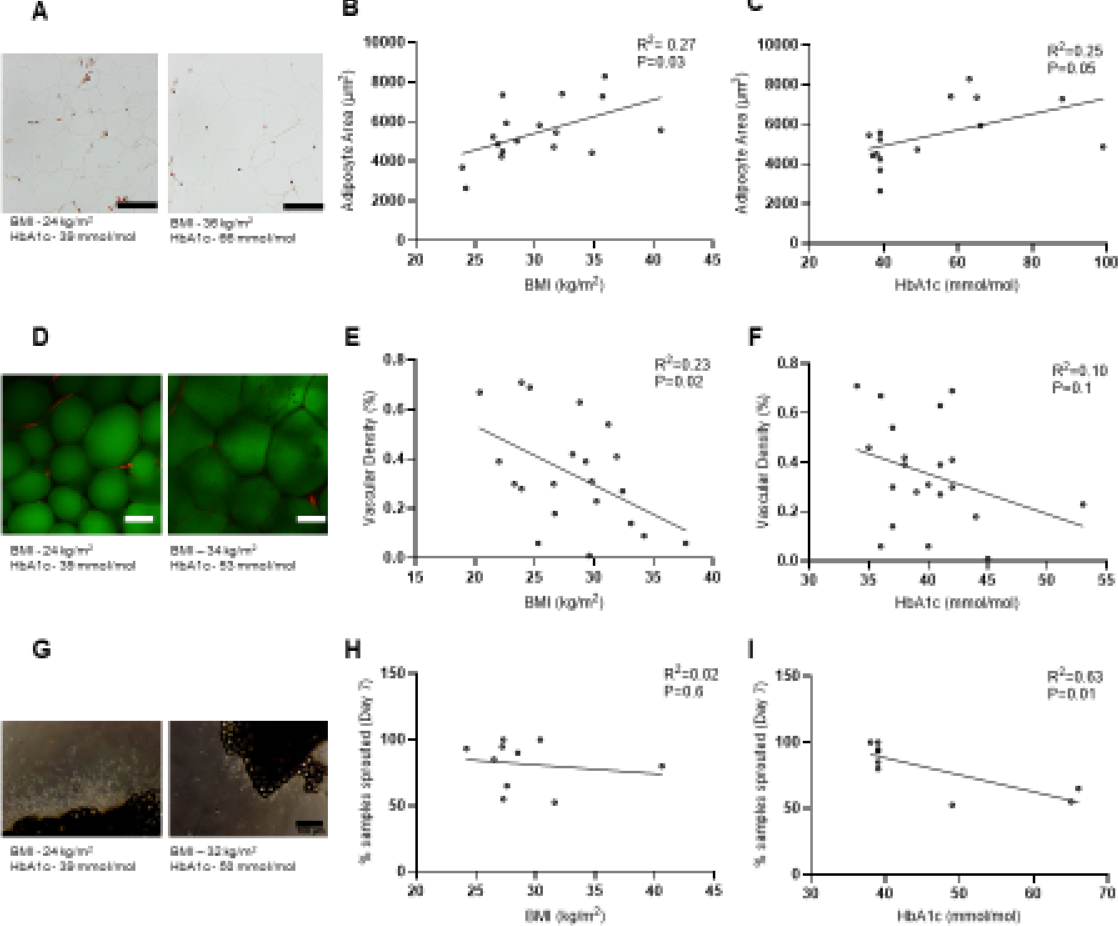
Deleterious remodelling of adipose tissue with increasing BMI and HbA1c. **A.** Representative images of hematoxylin and eosin (H & E)-stained white subcutaneous adipose tissue from patients with lower BMI and HbA1C and with higher BMI and HbA1c (Scale bar = 200µm). **B.** Correlation between BMI and adipocyte area (n =17 patients). **C.** Correlation between HbA1C and adipocyte area (n =17 patients). **D.** Representative images of Ulex Europaeus (Red) and LipidTox (Green) stained white subcutaneous adipose tissue from patients with lower BMI and HbA1C and with higher BMI and HbA1c (Scale bar = 100µm). **E.** Correlation between BMI and adipose vascularity (n= 21 patients). **F.** Correlation between HbA1C and adipose vascularity (n= 21 patients). **G.** Representative images of human white subcutaneous adipose tissue from patients with lower BMI and HbA1C and with higher BMI and HbA1c. (Scale bar = 100µm). **H.** Correlation between BMI and adipose neovascularisation (n= 10 patients). **I.** Correlation between HbA1C and adipose neovascularisation (n= 10 patients). Data points are individual patients. earsons’ correlation coefficients (*r*) were calculated to assess the degree of relation between BMI and Hba1C and various fat markers. p<0.05 taken as statistically significant.

**Supplementary figure 6.**
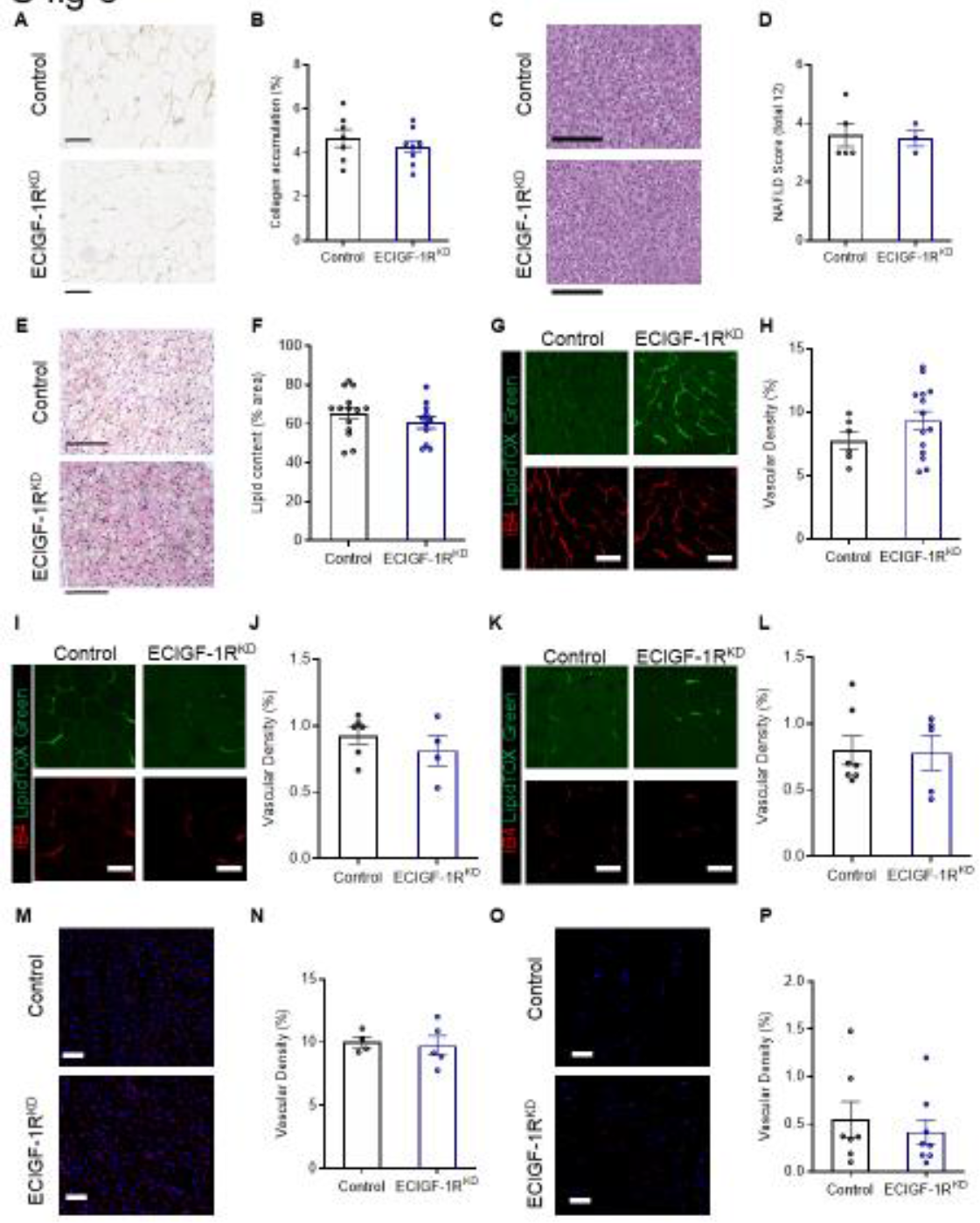
Histological characterisation of mice with reduced endothelial IGF-1R expression after 2 weeks of high fat diet. **A.** Representative images of picro sirius red stained white epididymal adipose tissue from 2-week HFD control and ECIGF-1R^KD^ mice (Scale bar = 200µm). **B.** Quantification of white epididymal adipose collagen deposition from 2-week HFD control and ECIGF-1R^KD^ mice (n =7&9). **C.** Representative images of Hematoxylin and eosin (H and E)-stained liver from 2-week HFD control and ECIGF-1R^KD^ mice (Scale bar = 200µm). **D.** Quantification of non-alcoholic fatty liver disease (NAFLD) from 2-week HFD control and ECIGF-1R^KD^ mice (n =5&3). **E.** Representative images of H and E-stained brown interscapular adipose tissue from 2-week HFD control and ECIGF-1R^KD^ mice (Scale bar = 100µm). **F.** Quantification of lipid content of interscapular brown adipose tissue from 2-week HFD control and ECIGF-1R^KD^ mice (n =14&12). **G.** Representative images of isolectin B4 (Red) and LipidTox (Green) stained brown interscapular adipose tissue from 2-week HFD control and ECIGF-1R^KD^ mice (Scale bar = 100µm). **H.** Quantification of vascularity in interscapular brown adipose tissue from 2-week HFD control and ECIGF-1R^KD^ mice (n =6&14). **I.** Representative images of isolectin B4 (Red) and LipidTox (Green) stained white subcutaneous adipose tissue from 2-week HFD control and ECIGF-1R^KD^ mice (Scale bar = 100µm). **J.** Quantification of vascularity in white subcutaneous adipose tissue from 2-week HFD control and ECIGF-1R^KD^ mice (n =6&4). **K.** Representative images of isolectin B4 (Red) and LipidTox (Green) stained white perinephric adipose tissue from 2-week HFD control and ECIGF-1R^KD^ mice (Scale bar = 100µm). **L.** Quantification of vascularity in white perinephric adipose tissue from 2-week HFD control and ECIGF-1R^KD^ mice (n =7&5). **M.** Representative images of isolectin B4 (Red) and DAPI (Blue) stained liver from 2-week HFD control and ECIGF-1R^KD^ mice (Scale bar = 50µm). **N.** Quantification of liver vascularisation from 2-week HFD control and ECIGF-1R^KD^ mice (n =4&5). **O.** Representative images of isolectin B4 (Red) and DAPI (Blue) stained muscle from 2-week HFD control and ECIGF-1R^KD^ mice (Scale bar = 50µm). **P.** Quantification of muscle vascularisation from 2-week HFD control and ECIGF-1R^KD^ mice (n =7&8). Data shown as mean ± SEM, Individual mice are shown as separate data points p<0.05 taken as being statistically significant using a student unpaired two tailed t-test and denoted as *.

**Supplementary figure 7.**
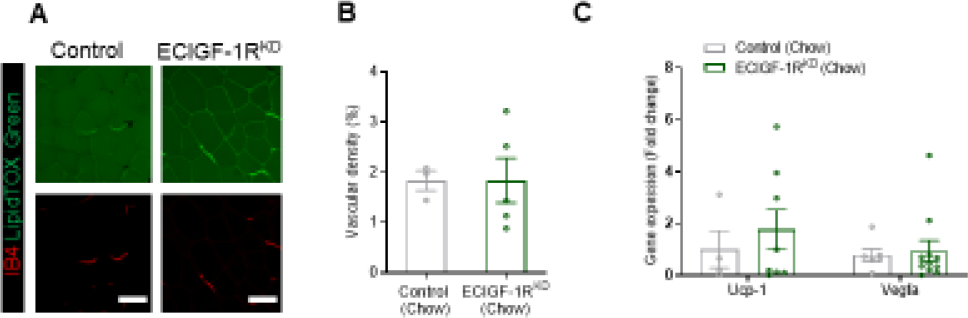
Characterisation of adipose tissue from chow fed mice with reduced endothelial IGF-1R expression. **A.** Representative images of isolectin B4 (Red) and LipidTox (Green) stained white epididymal adipose tissue from chow fed control and ECIGF-1R^KD^ mice (Scale bar = 100µm). **B.** Quantification of vascularity in white epididymal adipose tissue from chow fed control and ECIGF-1R^KD^ mice (n =3&5). **C.** Quantification of *Ucp-1* and *Vegfa* gene expression in white epididymal adipose tissue from chow fed control and ECIGF-1R^KD^ mice (n =4-11). Data shown as mean ± SEM, Individual mice are shown as separate datapoints p<0.05 taken as statistically significant using a student unpaired two tailed t-test and denoted as*.

**Supplementary figure 8.**
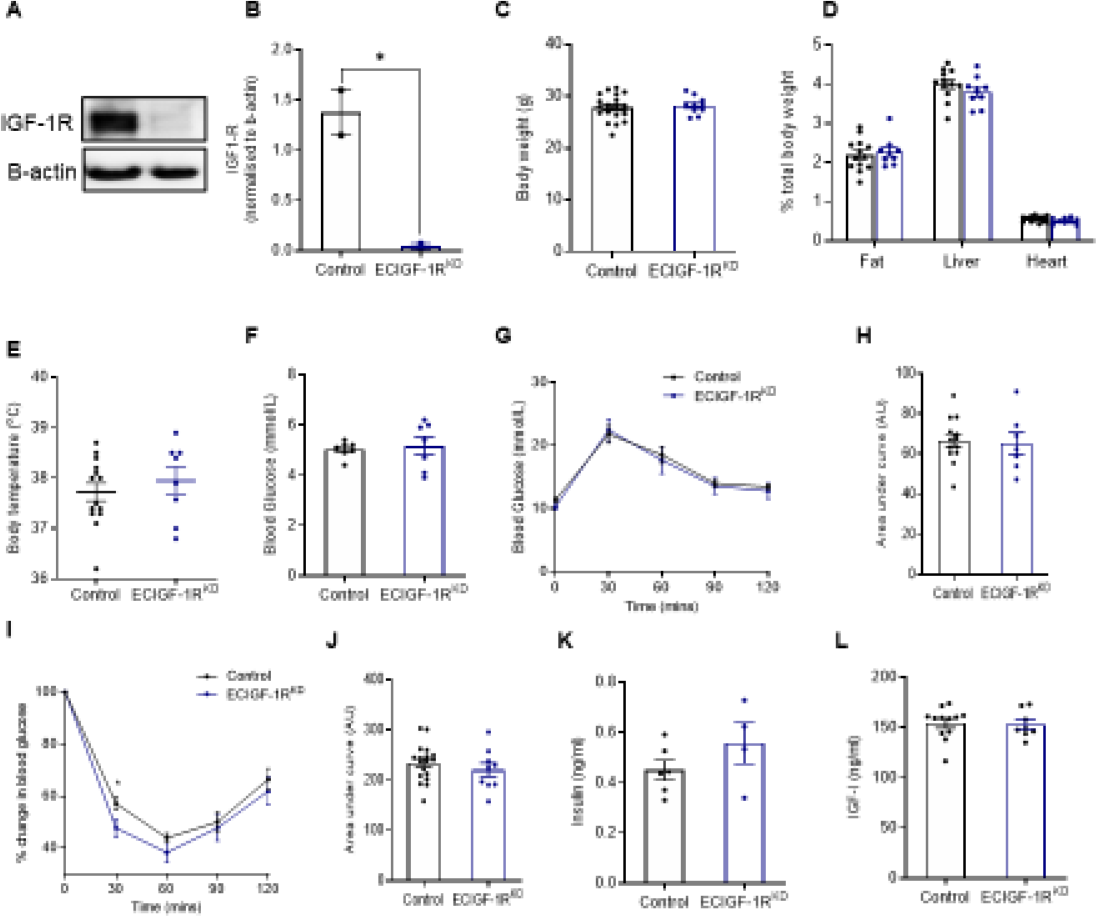
Metabolic characterisation of mice with reduced endothelial IGF-1R expression after 8-weeks high fat diet. **A.** Representative western blot of primary murine endothelial cell expression of IGF-1R from 8-week HFD fed control and ECIGF-1R^KD^ mice. **B.** Quantitation of primary murine endothelial cell expression of IGF-1R from 8-week HFD control and ECIGF-1R^KD^ mice (n=2&2). **C.** Body mass of 8-week HFD fed control and ECIGF-1R^KD^ mice (n=20&10). **D.** Wet organ weight of 8-week HFD control and ECIGF-1R^KD^ mice (n=13&9). **E.** Core body temperature of 8-week HFD fed control and ECIGF-1R^KD^ mice (n=14&8). **F.** Fasting blood glucose levels from 8-week HFD fed control and ECIGF-1R^KD^ mice (n=7&7) **G.** Glucose tolerance over time of 8-week HFD fed control and ECIGF-1R^KD^ mice (n=13&7). **H.** Area under the curve (AUC) analysis of glucose tolerance of 8-week HFD fed control and ECIGF-1R^KD^ mice (n=13&7). **I.** Insulin tolerance over time of 8-week HFD fed control and ECIGF-1R^KD^ mice (N=18&11). **J.** Area under the curve (AUC) analysis of insulin tolerance test of 8-week HFD fed control and ECIGF-1R^KD^ mice (n=18&11). **K.** Fasting plasma insulin levels from 8-week HFD fed control and ECIGF-1R^KD^ mice (n=6&4). **L.** Fasting plasma IGF-1 levels from 8-week HFD fed control and ECIGF-1R^KD^ mice (n=12&8). Data shown as mean ± SEM, individual mice are shown as separate datapoints p<0.05 taken as being statistically significant using a student unpaired two tailed t-test and denoted as *.

**Supplementary figure 9.**
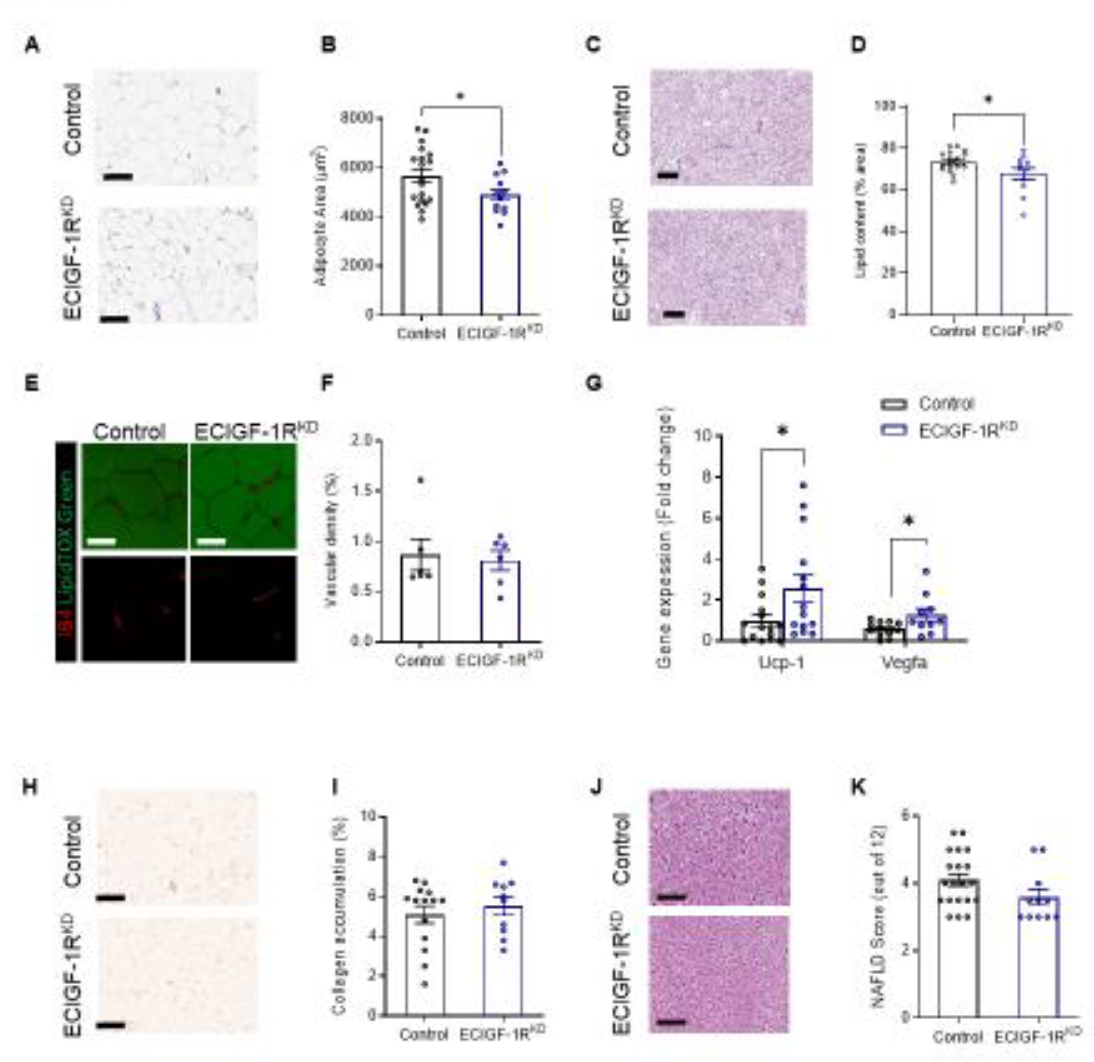
Histological characterisation of mice with reduced endothelial IGF-1R expression after 8-weeks of high fat diet. **A.** Representative images of Hematoxylin and eosin (H & E) stained white epididymal adipose tissue from 8-week HFD fed control and ECIGF-1R^KD^ mice (Scale bar = 200µm). **B.** Quantification of adipocyte size from 8-week HFD fed control and ECIGF-1R^KD^ mice (n= 12-18). **C.** Representative images of H & E stained brown interscapular adipose tissue from 8-week HFD fed control and ECIGF-1R^KD^ mice (Scale bar = 200µm). **D.** Quantification of lipid content of brown interscapular adipose tissue from 8-week HFD fed control and ECIGF-1R^KD^ mice (n =18&13). **E.** Representative images of isolectin B4 (Red) and LipidTox (Green) stained white epididymal adipose tissue from 8-week HFD control and ECIGF-1R^KD^ mice (Scale bar = 100µm). **F.** Quantification of white epididymal adipose tissue vascularisation from 8-week HFD fed control and ECIGF-1R^KD^ mice (n=6&6). **G.** Quantitation of white epididymal adipose gene expression of *Ucp-1* and *Vegfa* from 8-week HFD fed control and ECIGF-1R^KD^ mice. (n=12-15). **H.** Representative images of picro sirius red stained white adipose tissue from 8-week HFD fed control and ECIGF-1R^KD^ mice (Scale bar = 200µm). **I.** Quantification of white epididymal adipose collagen deposition from 8-week HFD fed control and ECIGF-1R^KD^ mice (n=14&10). **J.** Representative images of H and E-stained liver from 8-week HFD fed control and ECIGF-1R^KD^ mice (Scale bar = 200µm). **K.** Quantification of non-alcoholic fatty liver disease (NAFLD) from 8-week HFD fed control and ECIGF-1R^KD^ mice (n=20&11). Data shown as mean ± SEM, Individual mice are shown as separate datapoints p<0.05 taken as being statistically significant using a student unpaired two tailed t-test and denoted as *.

**Supplementary figure 10.**
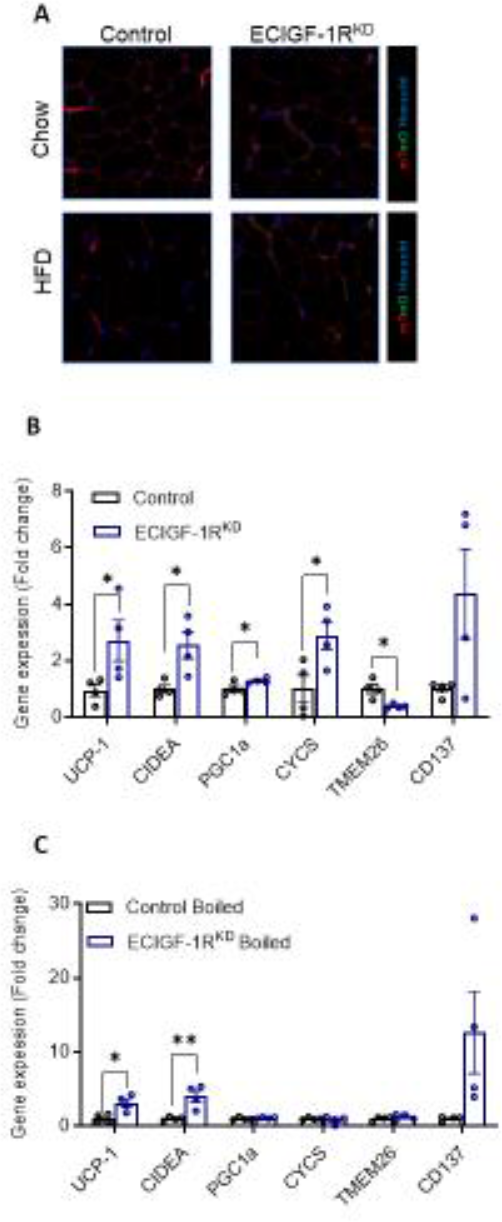
Reduction in murine endothelial IGF-1R expression alters the endothelial secretome and reveals a role for a small molecule in modulating adipocyte function. **A.** Following induction with tamoxifen, cells of endothelial cell lineage in the ECIGF-1R^KD^ fluoresce green using the mTmG system, with all other cells fluorescing red. All adipocytes from both genotypes appear red, confirming adipocytes are not from endothelial lineage. Quantitation of human primary adipocyte gene expression after 24hr treatment with conditioned media from primary murine endothelial cell isolated from 2-week HFD control and ECIGF-1R^KD^ mice. **B.** Quantitation of human primary adipocyte gene expression after treatment with boiled conditioned media from primary murine endothelial cells isolated from 2-week HFD fed control and ECIGF-1R^KD^ mice (n=4&4). Data shown as mean ± SEM, individual mice are shown as separate datapoints p<0.05 taken as being statistically significant using a student unpaired two tailed t-test and denoted as *.

**Supplementary figure 11.**
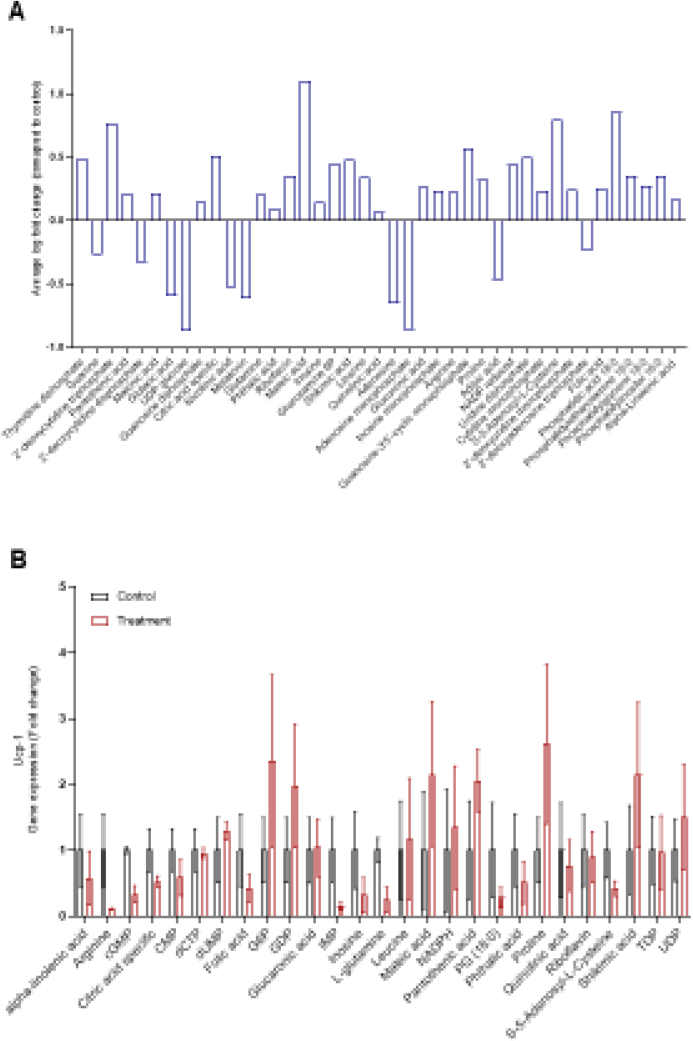
Mice with reduced endothelial IGF-1R expression after 2-weeks high fat diet have an altered endothelial small molecule secretome. **A.** Small molecule analysis of the aqueous and lipid fractions of conditioned media from primary murine endothelial cell from 2-week HFD fed control and ECIGF-1R^KD^ mice (n=4&4 per genotype). **B.** Quantitation of 3T3-L1 adipocyte gene expression of Ucp-1 after upregulated metabolite stimulation (n=3-5 per treatment group). Data shown as mean ± SEM, n is an experimental replicates p<0.05 taken as statistically significant using student t-test and denoted as *.

**Supplementary figure 12.**
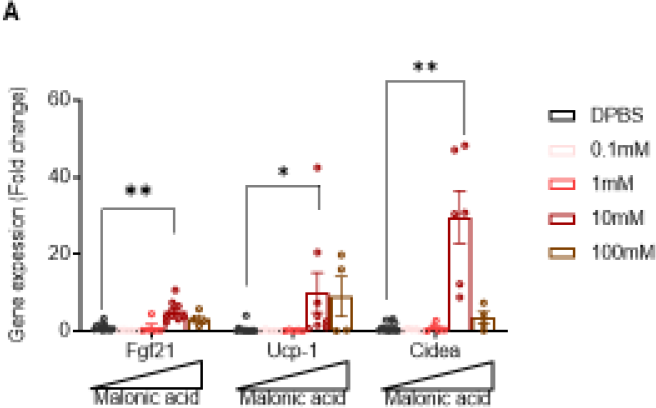
Malonic acid dose response. **A.** Quantification of gene expression of 3T3-L1 adipocytes treated with varying doses of malonic acid for 24hrs (n =5-14 per treatment group). Data shown as mean ± SEM, n is experimental replicates p<0.05 taken as being statistically significant using a one-way ANO A and denoted as * (p≤. and is denoted as **).

**Supplementary figure 13.**
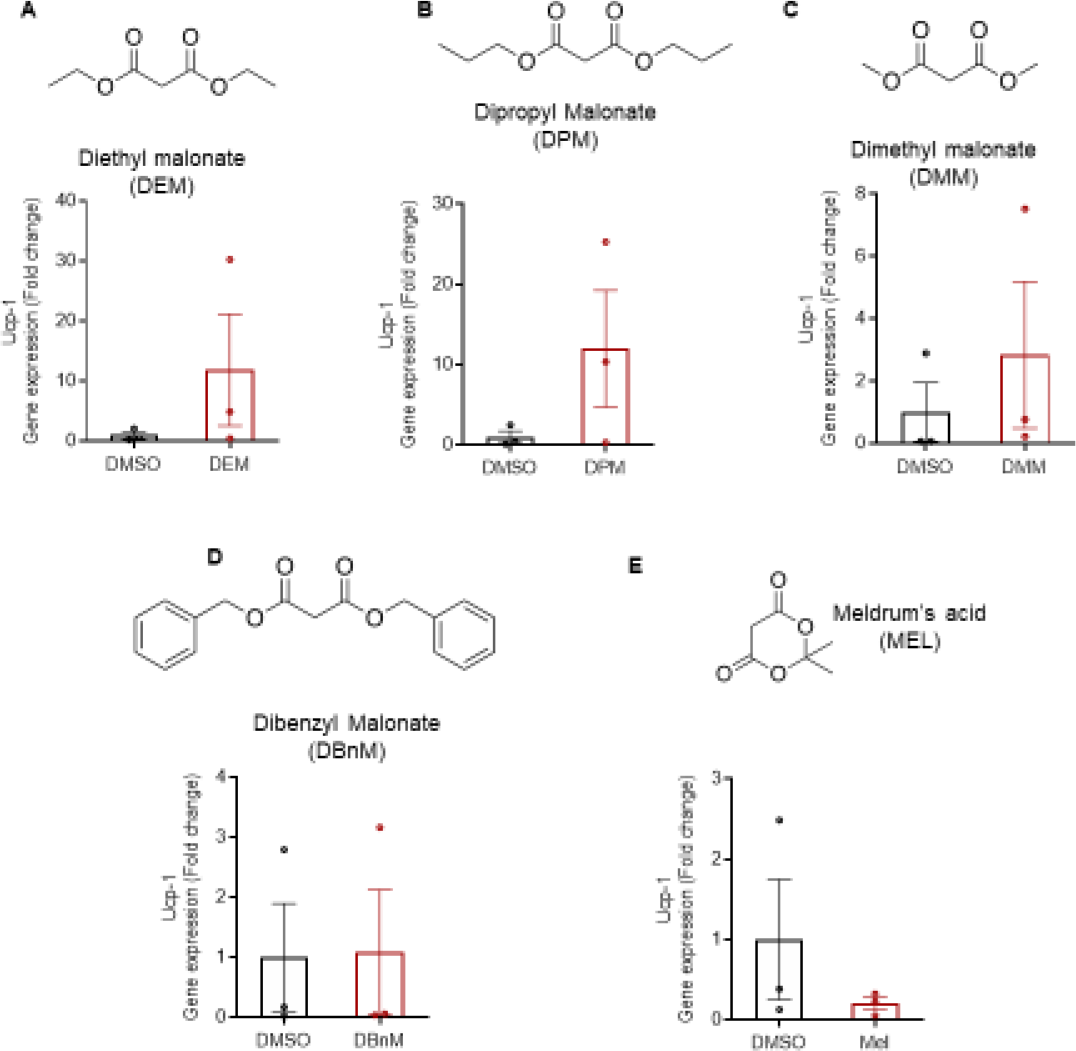
Screening of malonic acid pro-drugs in 3T3**-L1** adipocytes. **A.** Quantification of *Ucp-1* gene expression for 24hrs treatment with 10mM Diethyl malonate (DEM) (Chemical structure shown above). (n =3&3 per treatment group). **B.** Quantification of *Ucp-1* gene expression for 24hrs treatment with 10mM Dipropyl Malonate (DPM) (Chemical structure shown above). (n =3&3 per treatment group). **C.** Quantification of *Ucp-1* gene expression for 24hrs treatment with 10mM Dimethyl malonate (DMM) (Chemical structure shown above). (n =3&3 per treatment group). **D.** Quantification of *Ucp-1* gene expression for 24hrs treatment with 10mM Dibenzyl Malonate (DBnM) (Chemical structure shown above). (n =3&3 per treatment group). **E.** Quantification of *Ucp-1* gene e pression for 24hrs treatment with mM meldrum’s acid (MEL) (Chemical structure shown above). (n =3&3 per treatment group). Data shown as mean ± SEM, n is experimental replicates p<0.05 taken as statistically significant using an unpaired two sided student t-test and denoted as *.

